# Prolonged metaphase II arrest weakens Aurora B/C- dependent error correction in mouse oocytes

**DOI:** 10.1101/2023.10.11.561823

**Authors:** Antoine Langeoire, Alison Kem-Seng, Damien Cladière, Katja Wassmann, Eulalie Buffin

## Abstract

Chromosome segregation during meiosis is highly error-prone in mammalian oocytes. The mechanisms controlling chromosome attachments and the Spindle Assembly Checkpoint (SAC) have been extensively studied in meiosis I, but our knowledge of these mechanisms during meiosis II is rather limited. Although mammalian oocytes arrest in metaphase II for an extended period awaiting fertilization, some misattached chromosomes may persist. This suggests that the mechanism correcting misattachments is not fully functional during the arrest. In this study, we investigated whether low inter-kinetochore tension, which characterizes incorrect attachments, can be detected by Aurora B/C-dependent error correction in meiosis II. We found that low tension in early metaphase II does indeed mediate microtubule detachment by Aurora B/C and consequently, anaphase II delay through SAC activation. Surprisingly, we also found that during prolonged metaphase II arrest, Aurora B/C activity is no longer sufficient to detach low-tension attachments, concomitantly with high accumulation of PP2A at kinetochores. As a result, the SAC is not activated and sister chromatids segregate in anaphase II without delay even in the presence of low tension. Hence, during the prolonged metaphase II arrest to await fertilization, oocytes become unable to discriminate between correct and incorrect attachments and may allow errors to persist.

## INTRODUCTION

The formation of haploid gametes, which is essential for sexual reproduction, requires two consecutive meiotic divisions, with no S-phase in between. First, homologous chromosomes are segregated in meiosis I, followed by meiosis II with the segregation of sister chromatids. Meiosis must be carried out without any error to produce embryos with the correct ploidy after fertilization. Aneuploidy in the embryo is mainly due to mis-segregation events in the oocyte and is the leading cause of miscarriage, infertility and birth defects in humans^1–3^. Why female meiosis is highly error-prone is still a matter of debate, and possible explanations have been discussed in recent reviews^3–6^.

Mammalian oocytes enter meiosis after premeiotic S-phase. During prophase I, homologous chromosomes recombine and are held together by the resulting chiasmata, forming a bivalent. Oocytes remain arrested in prophase I and only resume meiosis at sexual maturity following hormonal stimulation. Nuclear envelope breakdown, also known as GVBD (Germinal Vesicle Breakdown) is followed by the formation of a bipolar spindle. In mouse oocytes, spindle assembly takes 4-6 hours and results in the attachment of homologous chromosomes to opposite poles of the spindle, with sister kinetochores oriented to the same pole^7,8^. Because initial attachments are often incorrect, they must be destabilized to give the microtubules a chance to re-attach correctly. Aurora B and the meiosis-specific Aurora C kinase are the enzymatic subunits of the Chromosomal Passenger Complex (CPC), which localizes at the centromere and interchromatid axis in meiosis I and promotes microtubule turnover at the kinetochore^9–13^. Thus, in meiosis I, bipolar attachments of homologous chromosomes are achieved after multiple rounds of Aurora B/C-dependent error correction^4,14^.

In mitosis, it was shown that the kinetochore-microtubule (KT-MT) interaction is controlled by the phosphorylation state of the NDC80 subunit Hec1, which anchors the microtubule plus ends to the kinetochore. KT-MT interactions are destabilized by Aurora B phosphorylation of Hec1 and stabilized by PP2A-B56-mediated dephosphorylation of Hec1^15–20^. The balance between Aurora B and PP2A-B56 activities is thought to depend on the tension that is generated between sister kinetochores when the opposing forces exerted by microtubules are in equilibrium. Inter-kinetochore (inter-KT) tension refers to the stretching of the centromeric region that has been observed when microtubules pull on kinetochores, in human or yeast mitotic cells^21,22^. The spatial separation model proposes that the inter-KT tension separates centromeric Aurora B from the kinetochore and reduces its ability to phosphorylate Hec1 and to destabilized KT-MT interaction. This model explains, at least in part, how Aurora B selectively destabilizes incorrect attachments, which are characterized by unbalanced pulling forces on sister kinetochores^4,23–25^.

In meiosis I, since sister kinetochores are mono-oriented and pulled in the same direction, tension is not generated between the sister kinetochores as in mitosis or meiosis II, but rather within kinetochores. As we have previously shown in mouse oocytes, reducing the forces pulling on kinetochores in metaphase I induces Hec1 phosphorylation by Aurora B/C, and Spindle Assembly Checkpoint (SAC) activation as a consequence of microtubule detachment^26^. Thus, despite mono-orientation of sister kinetochores, tension applied to paired kinetochores controls attachment correction in meiosis I. However, Aurora B/C persists in the vicinity of kinetochores even after biorientation of the bivalent, potentially preventing attachment stabilization. It has been proposed that attachment stabilization is mainly mediated by the progressive accumulation of PP2A-B56 phosphatase at kinetochores during meiosis, shifting the balance from Aurora B/C-dependent phosphorylation to dephosphorylation by PP2A-B56^27^.

After homologous chromosome segregation in anaphase I, followed by first polar body extrusion, oocytes immediately enter meiosis II. The transition from meiosis I to meiosis II, called interkinesis, is characterized by partial chromosome decondensation and the absence of a bipolar spindle. After interkinesis, spindle assembly in mouse oocyte meiosis II takes place within 1 hour, and is thus strikingly faster than in meiosis I^28,29^. In vertebrate oocytes, including mammals, meiosis arrests again at metaphase II, with sister chromatids attached in a bipolar fashion, in contrast to meiosis I^30^. This cell cycle arrest, maintained by CSF (Cytostatic Factor) activity, persists until fertilization takes place. Sperm entry triggers anaphase II with sister chromatid segregation, second polar body extrusion, and eventually zygote formation^31^(Figure 1A).

**Figure 1.**
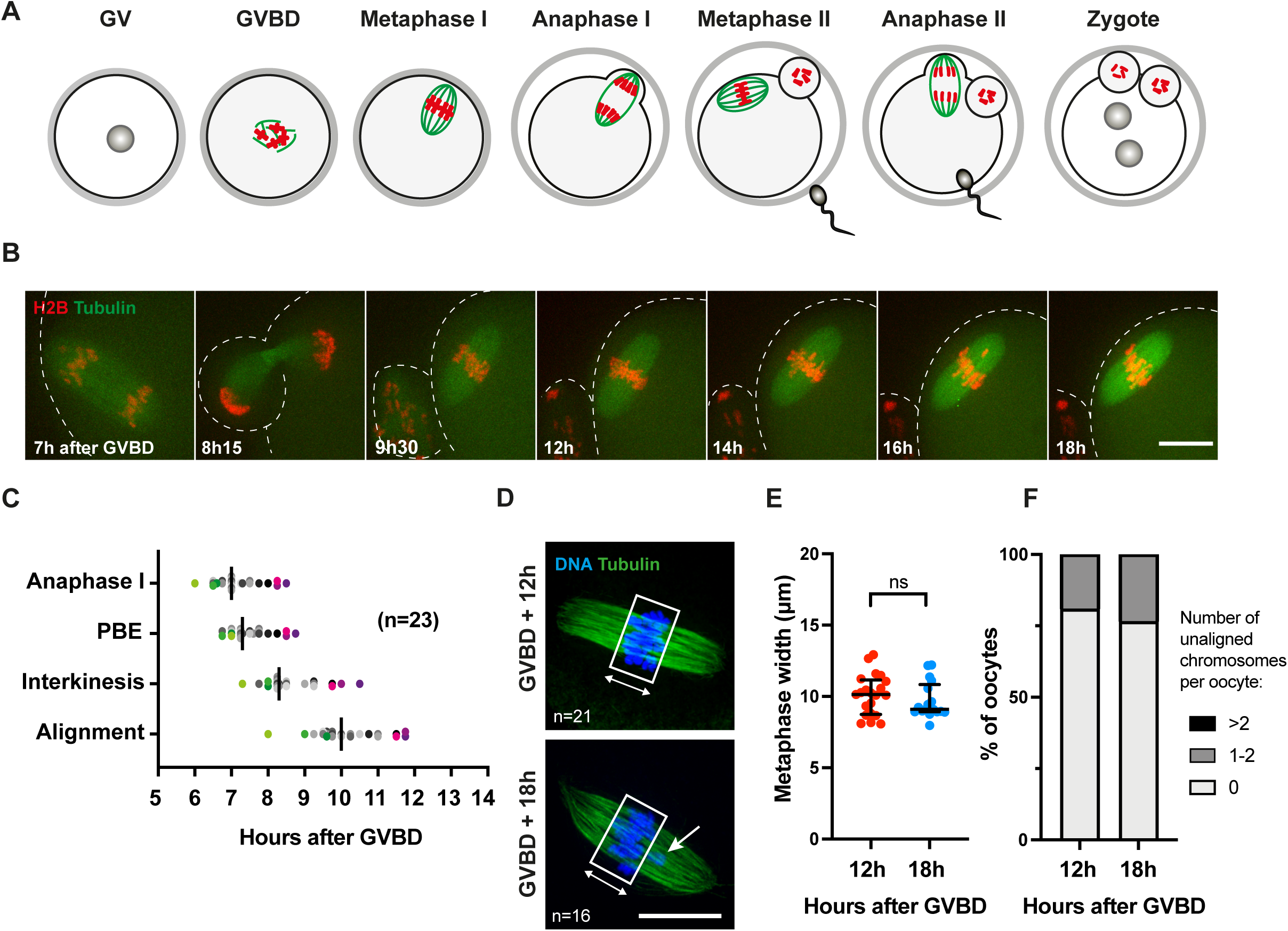
Unaligned chromosomes persist in metaphase II after *in vitro* maturation. (A) Scheme of meiotic maturation. Oocytes are arrested in prophase I (GV: germinal vesicle stage) and undergo GVBD (Germinal Vesicle Breakdown) upon hormonal stimulation. In meiosis I, homologous chromosomes (red) bi-orient to the spindle (green) and align in metaphase I before segregating in anaphase I. After extrusion of the first polar body, sister chromatids bi-orient and align in metaphase II. Oocytes arrest in metaphase II until fertilization. Sperm entry induces sister chromatid segregation in anaphase II, second polar body extrusion, and zygote formation. Female and male nuclei will fuse during the first mitotic division. (B) Prophase I arrested oocytes were injected with H2B-RFP and Tubulin-GFP mRNA, released and synchronised at GVBD (Germinal Vesicle Breakdown). *In vitro* maturation was observed by time-lapse microscopy from anaphase I to metaphase II. Hours after GVBD are indicated. Scale bar: 20 µm. (C) Timing of anaphase I, polar body extrusion (PBE), interkinesis, and first alignment of chromosomes in 23 oocytes analysed as in B. Each dot represents an oocyte. The green and purple dots correspond respectively to the three earliest and the three latest oocytes doing anaphase I. Median is indicated. n: number of oocytes. (D) *In vitro* matured oocytes were fixed in meiosis II at 12 h or 18 h after GVBD, and stained with DAPI (blue) and anti-Tubulin antibody (green). The rectangle defines the metaphase plate, and the double-headed arrow indicates how the width of the metaphase plate was measured in (E). Arrow indicates unaligned chromosome as quantified in (F). Scale bar: 20 µm. n is the number of oocytes analyzed from at least 3 independent experiments for each condition. (E) The width of the metaphase plate was measured in each oocyte from (D). Median and interquartile range are indicated. p values have been calculated by using Mann-Whitney’s test (*p<0.05, ns: not significant). (F) The percentage of oocytes containing 0, 1-2, or more than 2 unaligned chromosomes, was quantified in each oocyte from (D). Note that more than 2 unaligned chromosomes were never observed under our conditions. Chromosomes were considered unaligned if they were clearly outside the metaphase plate as shown in (D).

Since mammalian oocytes are arrested in metaphase II until fertilization, they should have more time to correct faulty chromosome attachments. However, in mouse metaphase II oocytes, 20% of attachments are merotelic or lateral, suggesting that detection and correction of incorrect attachments in meiosis II is compromised^28^. In addition, meiosis II has to deal with segregation errors carried over from meiosis I. In mouse oocytes, single sister chromatids resulting from precocious separation in meiosis I can escape error correction and SAC activation in meiosis II^32^. Therefore, it is essential to better understand how correct attachments are achieved during meiosis II, and why errors persist and are not always recognized.

Here, to investigate the efficiency of the Aurora B/C-dependent error correction pathway in meiosis II, we asked whether low-tension attachments can be detected and corrected in meiosis II. Unexpectedly, we found that after prolonged arrest in metaphase II, low tension does not promote Aurora B/C-dependent microtubule detachment, or SAC activation, in contrast to reducing tension at an earlier stage. We conclude that when oocytes are kept arrested in metaphase II to await fertilization, attachments, even incorrect ones, are stabilized by PP2A-B56 accumulating at the kinetochore and counteracting Aurora B/C, regardless of the level of tension. This may explain the persistence of misaligned chromosomes in metaphase II oocytes.

## RESULTS

### Unaligned chromosomes persist in metaphase II after *in vitro* maturation

First, to define when exactly oocytes are in metaphase II after *in vitro* maturation, we characterized spindle reformation and chromosome alignment after anaphase I by time-lapse microscopy in mouse oocytes expressing H2B-RFP and Tubulin-GFP (Figure 1B). In our strain background, chromosomes segregated in anaphase I approximately 7 h after GVBD. After polar body extrusion, chromosomes appeared partially decondensed but remained close together, even though the spindle had not yet reassembled. Interkinesis lasted for about 1 h. Then, the chromosomes individualized, and the metaphase II spindle rapidly reassembled. Chromosomes began to align at the equator around 10 h after GVBD. Visualization of a metaphase plate defined the transition from prometaphase II to metaphase II, even though chromosomes continued to oscillate throughout the metaphase II arrest (Figure 1B-C and Video 1).

In *in vitro* matured oocytes fixed at 12 h or 18 h after GVBD, a metaphase plate of similar width was clearly visible (Figures 1D and 1E). However, although most chromosomes were correctly aligned, we observed that 20% of the oocytes contained one or two chromosomes outside the metaphase plate, even at 18 h after GVBD (Figure 1F). This is consistent with previous results reporting that incorrect attachments persist during metaphase II arrest^28^ and suggests that attachment errors may escape correction. Thus, we compared the efficiency of SAC and Aurora B/C-dependent error correction at 12 h after GVBD, shortly after chromosome alignment (early metaphase II), and at 18 h after GVBD (late metaphase II) to address whether prolonged arrest affects the detection of attachment errors.

### The SAC is functional throughout metaphase II

We first asked whether the SAC remains functional in early and late metaphase II and whether it is able to block anaphase II onset upon nocodazole-induced missing attachments^33^. To assess SAC activity after *in vitro* maturation, we monitored APC/C-dependent degradation of exogenously expressed Securin-YFP. *In vivo*, anaphase II is induced by fertilization, which can be mimicked *in vitro*, by activating oocytes with strontium treatment (Figure 2A). In the absence of nocodazole, Securin-YFP was rapidly degraded after addition of strontium. In contrast, Securin-YFP remained constant after addition of nocodazole either in early or late metaphase II, showing that the SAC is functional and can be efficiently activated upon gross loss of kinetochore attachments (Figures 2B, S1 and videos 2 and 3). Accordingly, the SAC protein Mad2, which was almost completely absent from kinetochores in control oocytes, was recruited to unattached kinetochores after depolymerization of microtubules with nocodazole in both early and late metaphase II, indicating that the SAC was well functional at both time points (Figures 2C and 2D). All together, these results show that the SAC retains its functionality in early and late metaphase II after *in vitro* maturation, and could theoretically be activated if low-tension attachments are released by the destabilizing activity of Aurora B/C.

**Figure 2.**
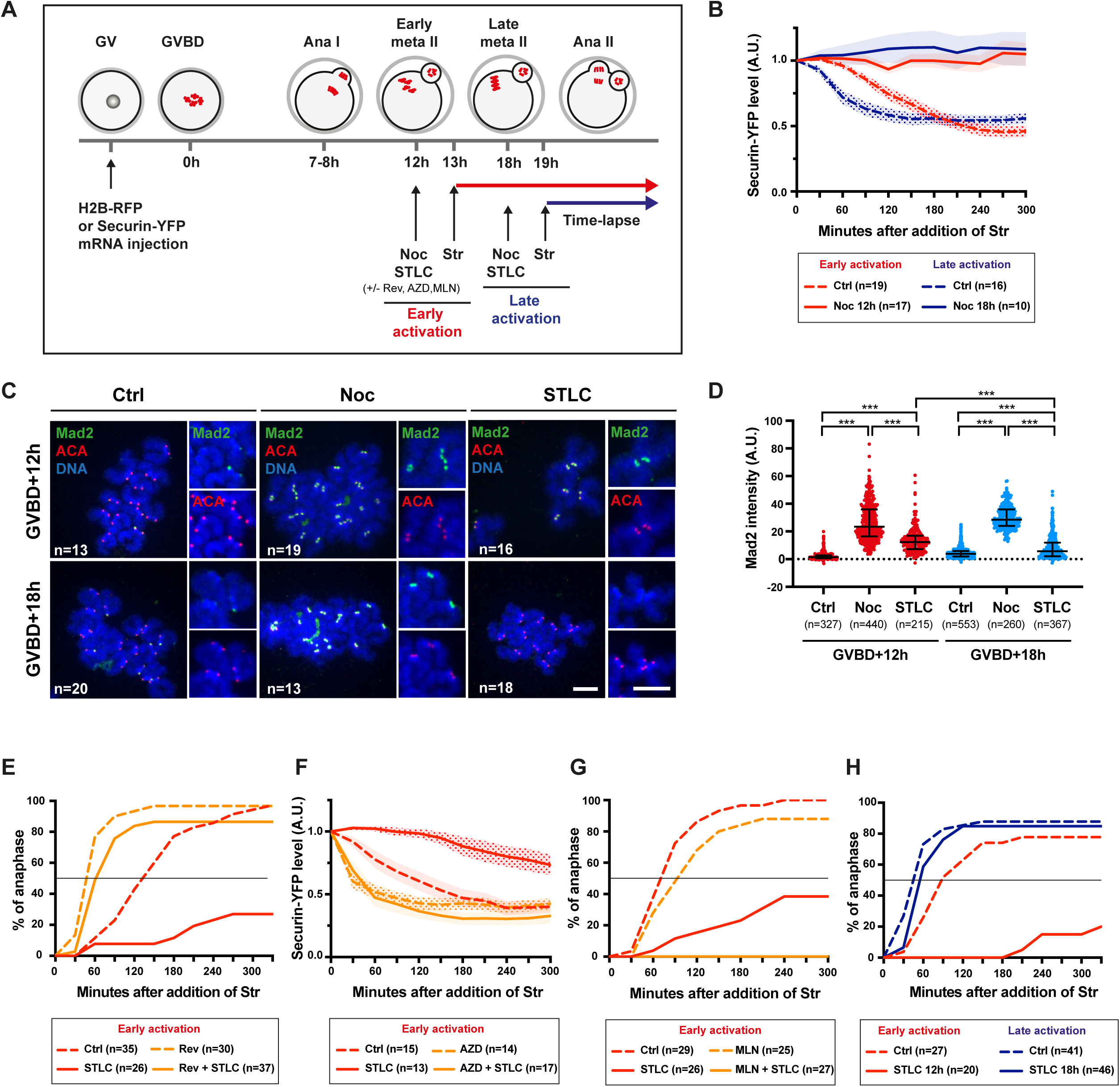
Low tension in early metaphase II but not late metaphase II induces SAC activation in an Aurora B/C-dependent way. (A) Scheme of the experimental settings in the panels (B) and (E-H) below: prophase I arrested oocytes were injected with the indicated mRNAs, released and synchronised at GVBD (Germinal Vesicle Breakdown). Meiosis II oocytes were submitted to drug treatment as indicated, at 12h or 18h after GVBD. Then, oocytes were activated by addition of strontium (Str) and their ability to progress into anaphase II was analysed by time-lapse imaging. (B) Oocytes injected with Securin-YFP mRNA were left untreated (Ctrl) or treated with nocodazole 0,4 µM (Noc) for 1 h, at GVBD+12h (red curves) or GVBD+18h (blue curves). After addition of strontium, the Securin-YFP levels were quantified in each oocyte for each time point of the time-lapse. n: number of oocytes from at least 3 experiments. Curves correspond to mean values, shading to SEM. (C) *In vitro* matured meiosis II oocytes were left untreated (Ctrl) or treated with Nocodazole 0,4 µM (Noc) or STLC 1 µM for 1 h, at GVBD+12h or GVBD+18h. Chromosomes were spread and stained with ACA (red), anti-Mad2 antibody (green) and DAPI (blue). Smaller images correspond to magnifications of chromosomes. Scale bars: 10 µm. n: number of oocytes. (D) The mean intensity of Mad2 signal per centromere from (C) has been quantified and normalized to the mean intensity of ACA (AU: arbitrary unit). The values from 3 independent experiments are shown and median and interquartile range are indicated. p values have been calculated by using Mann-Whitney’s test (*** p<0.0001). n: number of centromeres. (E) Oocytes injected with H2B-RFP mRNA were left untreated (Ctrl) or treated with STLC 1 µM alone (STLC), reversine 0,5 µM alone (Rev), or with both STLC and reversine (STLC+Rev), for 1 h at GVBD+12h. The percentage of oocytes undergoing anaphase II relative to the time after addition of strontium was determined by time-lapse microscopy. The straight horizontal line represents 50% of the oocytes that have entered anaphase II. n: number of oocytes. (F) Oocytes injected with Securin-YFP were left untreated (Ctrl) or treated with STLC 1 µM alone (STLC), AZD1152 0,5 µM alone (AZD), or with both STLC and AZD1152 (STLC+AZD), for 1 h at GVBD+12h. After addition of strontium, the level of Securin-YFP was quantified in each oocyte for each time point of the time-lapse. n: number of oocytes from at least 3 experiments. Curves correspond to mean values, shading to SEM. (G) Oocytes incubated with SirDNA were left untreated (Ctrl) or treated with STLC 1 µM alone (STLC), MLN8237 1 µM alone (MLN), or with both STLC and MLN8837 (STLC+MLN), for 1 h at GVBD+12h. The percentage of oocytes undergoing anaphase II relative to the time after addition of strontium was determined by time-lapse microscopy. The straight horizontal line represents 50% of the oocytes that have entered anaphase II. n: number of oocytes. (H) Oocytes injected with H2B-RFP mRNA were left untreated (Ctrl) or treated with STLC 1 µM (STLC) for 1 h, at GVBD+12h (red curves) or GVBD+18h (blue curves). The percentage of oocytes undergoing anaphase II relative to the time after addition of strontium was determined by time-lapse microscopy. The straight horizontal line represents 50% of the oocytes that have entered anaphase. n: number of oocytes.

### Low tension in early metaphase II induces SAC activation in an Aurora B/C-dependent way

Since the SAC is functional in early and late metaphase II, we hypothesized that the tension-sensing mechanism itself is inefficient to sense and thus correct low-tension attachments in metaphase II. Therefore, we tested first CPC functionality after *in vitro* maturation, in early metaphase II. STLC, an inhibitor of the kinesin Eg5, was used to provoke low-tension attachments in early metaphase II^34^. By promoting the pole-separating forces required for spindle bipolarization, Eg5 contributes to the pulling forces on kinetochores^7,35^. In meiosis I, Eg5 inhibition by STLC induced shorter spindles and reduced the distance between pairs of sister kinetochores, resulting in activation of Aurora B/C-dependent error correction^26^. As expected, STLC also reduced spindle length and inter-KT distance in early metaphase II, confirming the efficiency of Eg5 inhibition (Figures S2A-D). Inter KT-distance is commonly used to estimate the tension applied to sister kinetochore pairs^15,16^. Importantly, STLC does not affect overall attachment and chromosome alignment, as evidenced by super resolution images obtained with SIM (Video 4). To investigate whether low tension can induce microtubule detachment by the CPC and consequently, SAC activation, we asked whether anaphase II onset is delayed as expected when the SAC detects unattached kinetochores. Timing of anaphase II onset, after *in vitro* maturation, was determined by following sister chromatid segregation in H2B-RFP-expressing oocytes by live imaging (Figure 2A). When control oocytes were activated with strontium in early metaphase II (12 h after GVBD), 80 to 100 % of oocytes completed anaphase II and 50% of oocytes had already segregated sister chromatids within 2 h after strontium addition (Figures 2E, 2G and 2H: dotted red curves). Crucially, when STLC was added in early metaphase II, anaphase II was strongly delayed and less than 40% of STLC-treated oocytes segregated sister chromatids within 5 h after addition of strontium (Figures 2E, 2G and 2H: solid red curves, Figure S3A and Video 5). Accordingly, the SAC protein Mad2 was recruited to kinetochores when oocytes were treated with STLC in early metaphase II, confirming that the SAC was activated upon reduced tension (Figures 2C and 2D). Moreover, after inhibition of the essential SAC protein Mps1 with the drug reversine, STLC-treated oocytes in early metaphase II were now able to enter anaphase II after strontium addition, due to override of the SAC. Sister chromatids segregated even earlier than in control oocytes (less than 1 h for 50% of oocytes), supporting that the SAC was still active and was delaying anaphase II entry in early metaphase II control oocytes (Figure 2E: solid orange curve, Figure S3A and Video 5). Thus, in early metaphase II, the SAC is well activated after inducing low tension, suggesting that low-tension attachments can be detected and destabilized.

Next, we asked whether the anaphase delay observed after artificially reducing tension in early metaphase II, requires CPC activity. We chose to follow Securin-YFP degradation as an indicator of anaphase onset, because Aurora B/C inhibition perturbs sister chromatid segregation, making anaphase II onset more difficult to visualize. In STLC-treated early metaphase II oocytes, Securin-YFP degradation was delayed after strontium addition compared to control oocytes (Figure 2F: red curves and Figure S3B). After inhibition of Aurora B/C with AZD1152, STLC-treated oocytes in early metaphase II were able to degrade Securin-YFP without delay, indicating that SAC was satisfied and did not delay APC/C activation (Figure 2F: solid orange curve, Figure S3B and Video 6).

Because the dose of AZD1152 used in this experiment has been reported to also inhibit Aurora A^36^, we verified that Aurora A activity did not contribute to the anaphase II delay when STLC was added in early metaphase II. After inhibition of Aurora A with MLN8237, STLC-treated oocytes in early metaphase II were unable to segregate sister chromatids after addition of strontium (Figure 2G: solid orange curve, Figure S3C and video 7). Thus, artificially reducing tension in early metaphase II prevented anaphase II from occurring only when Aurora B/C was active, but Aurora A was not required. This indicates that in early metaphase II, low-tension attachments promote Aurora B/C-dependent error correction. As a result, the SAC was activated and blocked anaphase II onset, similar to meiosis I, as we have shown previously^26^.

### Low tension in late metaphase II does not induce SAC activation

For up to now, our data show that in early metaphase II, artificial reduction of inter-KT tension can be efficiently detected by Aurora B/C and can induce a SAC-dependent delay of anaphase II onset. However, few unaligned chromosomes can be observed in 20% of oocytes in late metaphase II (Fig 1F). Thus, we hypothesized that error correction was less efficient in late metaphase II.

To address this issue, we treated *in vitro* matured oocytes in late metaphase II with STLC to determine whether low tension induces a SAC-dependent anaphase II delay after activation with strontium (Figure 2A). As expected, STLC reduced spindle length and inter-KT distance in late metaphase II oocytes, without affecting chromosome attachment, confirming the efficiency of Eg5 inhibition (Figure S2E-H and video 8). When STLC was added in late metaphase II, 80% of oocytes were able to complete anaphase II and 50% of oocytes segregated sister chromatids within 1h after strontium addition, similar to untreated control oocytes activated in late metaphase II (Figure 2H: blue curves, Figure S3D and Video 9). Anaphase II was therefore induced rapidly, even after STLC treatment in late metaphase II oocytes. Thus, low-tension attachments did not promote SAC activation in late metaphase II. Accordingly, the SAC protein Mad2 remained at low levels on most kinetochores when oocytes were treated with STLC in late metaphase II after *in vitro* maturation (Figures 2C and 2D). Importantly, a higher dose of STLC (2 µM instead of 1 µM) reduces spindle length and inter-KT distance slightly more (Figure S2E-H), but in 30% of oocytes, the spindle almost collapses and chromosomes may have lost their attachment (Figure S2E-F). In this case, anaphase II was slightly delayed: 50% of oocytes treated with 2 µM STLC segregated sister chromatids within 2 h instead of 1h in 50% of control oocytes. Importantly, 60% of the oocytes treated with 2 µM STLC were still able to enter anaphase II at the end of the movie, suggesting that the low tension is not efficiently detected (Figure S2I). Accordingly, 40% of oocytes treated with 2 µM STLC failed to complete anaphase II compared to 20% of control oocytes. The additional 20% of 2 µM STLC- treated oocytes that failed to complete anaphase II may correspond to collapsed spindle oocytes, in which the SAC is activated by unattached kinetochores. Our results indicate that although the SAC is functional in early and late metaphase II, it is activated by low tension in early metaphase II but not in late metaphase II, after *in vitro* maturation of oocytes. Thus, there would be a temporal window during which low tension could be detected and induce a SAC-dependent delay to repair faulty attachments. In late metaphase II, low tension would be tolerated and thus become unable to activate error correction and thus, the SAC.

### PP2A accumulates in late metaphase II, counteracting Aurora B/C and stabilizing low-tension attachments

These latter results suggest that Aurora B/C-dependent error correction is efficient enough to promote microtubule detachment in early metaphase II but not in late metaphase II. To confirm that low-tension attachments promote destabilization of KT-MT interactions in early but not late metaphase II, we investigated the concomitant phosphorylation of Hec1 on Serine 55, which is a substrate of Aurora B/C and serves as a read-out of MT destabilization and ongoing error correction in mitosis^16^. The levels of Hec1 phosphorylation were similar in early and late metaphase II control oocytes, suggesting that most attachments are already stabilized in early metaphase II, although some unaligned chromosomes persist (Figures 3A and 3B). It is reasonable to assume that unaligned chromosomes show higher Hec1 phosphorylation, however, we did not compare Hec1 phosphorylation between aligned and unaligned chromosomes here. Crucially, treating oocytes with STLC in early metaphase II increased Hec1 phosphorylation significantly more than in late metaphase II (Figures 3A and 3B), consistent with the observed SAC activation and anaphase delay.

**Figure 3.**
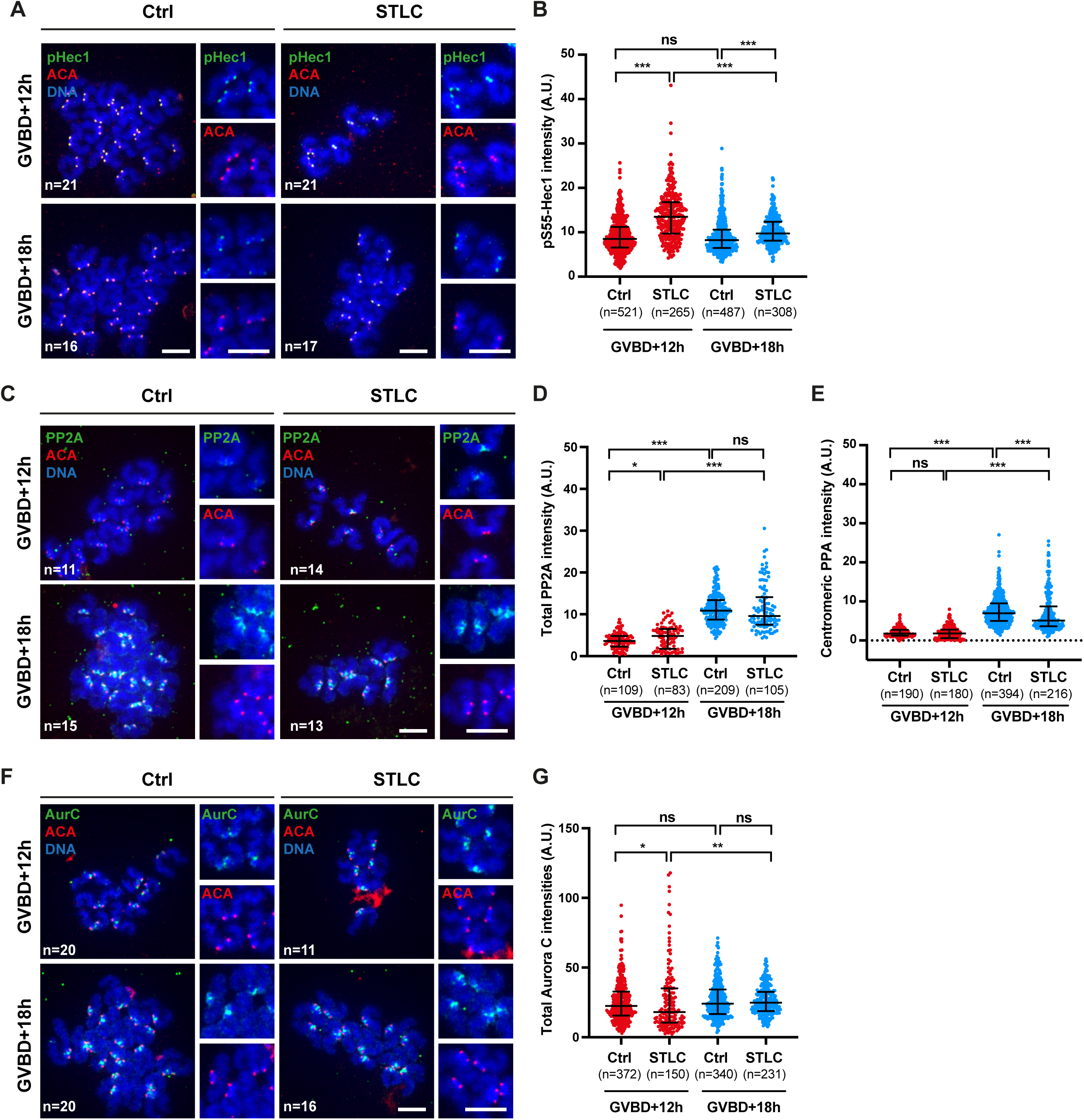
PP2A accumulates in late metaphase II, counteracting Aurora B/C and stabilizing low-tension attachments. (A) *In vitro* matured meiosis II oocytes at 12h or 18h after GVBD were treated with STLC at 1 µM for 1 h where indicated, and chromosomes were spread and stained with ACA (red), anti-pS55 Hec1 antibody (green) and DAPI (blue). Smaller images correspond to magnifications of chromosomes. Scale bar: 10 µm. n is the number of oocytes. (B) The mean intensity of pS55 Hec1 signal per centromere has been quantified and normalized to the mean intensity of ACA (AU: arbitrary unit). n is the number of centromeres analyzed in oocytes of (A) from 3 independent experiments. (C) *In vitro* matured meiosis II oocytes at 12h or 18h after GVBD were treated with STLC at 1 µM for 1 h where indicated, and chromosomes were spread and stained with ACA (red), anti-PP2A antibody (green) and DAPI (blue). Smaller images correspond to magnifications of chromosomes. Scale bar: 10 µm. n is the number of oocytes. (D) The mean intensity of total PP2A signal including kinetochores and between centromeres has been quantified and normalized to the mean intensity of ACA (AU: arbitrary unit). n is the number of centromere pairs analyzed in oocytes of (C) from 3 independent experiments. (E) The mean intensity of kinetochore PP2A signal has been quantified and normalized to the mean intensity of ACA (AU: arbitrary unit). n is the number of centromeres analyzed in oocytes of (C) from 3 independent experiments. (F) *In vitro* matured meiosis II oocytes at 12h or 18h after GVBD were treated with STLC at 1 µM for 1 h where indicated, and chromosomes were spread and stained with ACA (red), anti-Aurora C antibody (green) and DAPI (blue). Smaller images correspond to magnifications of chromosomes. Scale bar: 10 µm. n is the number of oocytes. (G) The mean intensity of total Aurora C signal including centromeres and between centromeres has been quantified and normalized to the mean intensity of ACA (AU: arbitrary unit). n is the number of centromere pairs analyzed in oocytes of (F) from 3 independent experiments. In all the graphs, median and interquartile range are indicated. p values have been calculated by using Mann-Whitney’s test (*** p<0.0001, **p< 0.001, *p<0.05, ns: not significant).

The level of Hec1 phosphorylation results from the balance between Aurora B/C kinase and PP2A- B56 phosphatase activities. Inefficient phosphorylation of Hec1 on Serine 55 in late metaphase II may be due to a shift in the balance towards higher PP2A-B56 activity, compared to early metaphase II. More PP2A-B56 activity may counteract Aurora B/C-dependent phosphorylation of Hec1 in late metaphase II, as already shown in meiosis I^27^. Hence, we compared PP2A levels in early and late metaphase II, after *in vitro* maturation. PP2A was localized at kinetochores around the centromeric ACA staining, and also between sister centromeres (Figure 3C), similar to Sgo2^37^. Indeed, levels of total PP2A were significantly higher in late than in early metaphase II, and only weakly affected by STLC at both metaphase II stages (Figures 3C and 3D). We also confirmed that the pool of PP2A located at kinetochores, which is most likely the one dephosphorylating Hec1, was increased in late metaphase II (Figure 3E). In contrast, levels of total Aurora C, located mainly between sister centromeres but also on some centromeres, remained constant between early and late metaphase II and were hardly affected by STLC (Figures 3F and 3G).

Based on these results, we propose that PP2A accumulates at kinetochores during metaphase II and antagonizes Aurora B/C, which remain constant. Thus, PP2A ultimately stabilizes kinetochore attachments in late metaphase II independently of low tension.

### Monopolar attachments can be corrected in early and late metaphase II through Aurora A activity

If low tension is not efficiently detected in late metaphase II, we expected that incorrect attachments induced at this stage would not be corrected. To test this hypothesis, we generated incorrect attachments by creating monopolar spindles after exposing early and late metaphase II oocytes to high doses of STLC. On these monopolar spindles, sister chromatids are attached to the same pole and are therefore under low tension. We asked whether these oocytes with a monopolar spindle had the ability to reform a bipolar spindle with correct attachments only in early metaphase II, but not in late metaphase II.

To determine whether these monopolar attachments can be corrected, oocytes were released from STLC treatment, to allow spindle bipolarization. After STLC removal, the time required for chromosomes to detach, reattach and align at the metaphase plate was measured by time-lapse microscopy, visualizing chromosomes with SirDNA (Figure 4A). Of note, similar experiments have been performed in mitosis to study the mechanisms underlying error correction^38^. The timing of chromosome realignment was used as a read-out for error correction efficiency. If error correction is functional, Aurora B/C should promote destabilization and correction of monopolar attachments, and chromosomes should realign correctly. We expected functional error correction in early but not late metaphase II, where high levels of PP2A should stabilize monopolar attachments.

**Figure 4.**
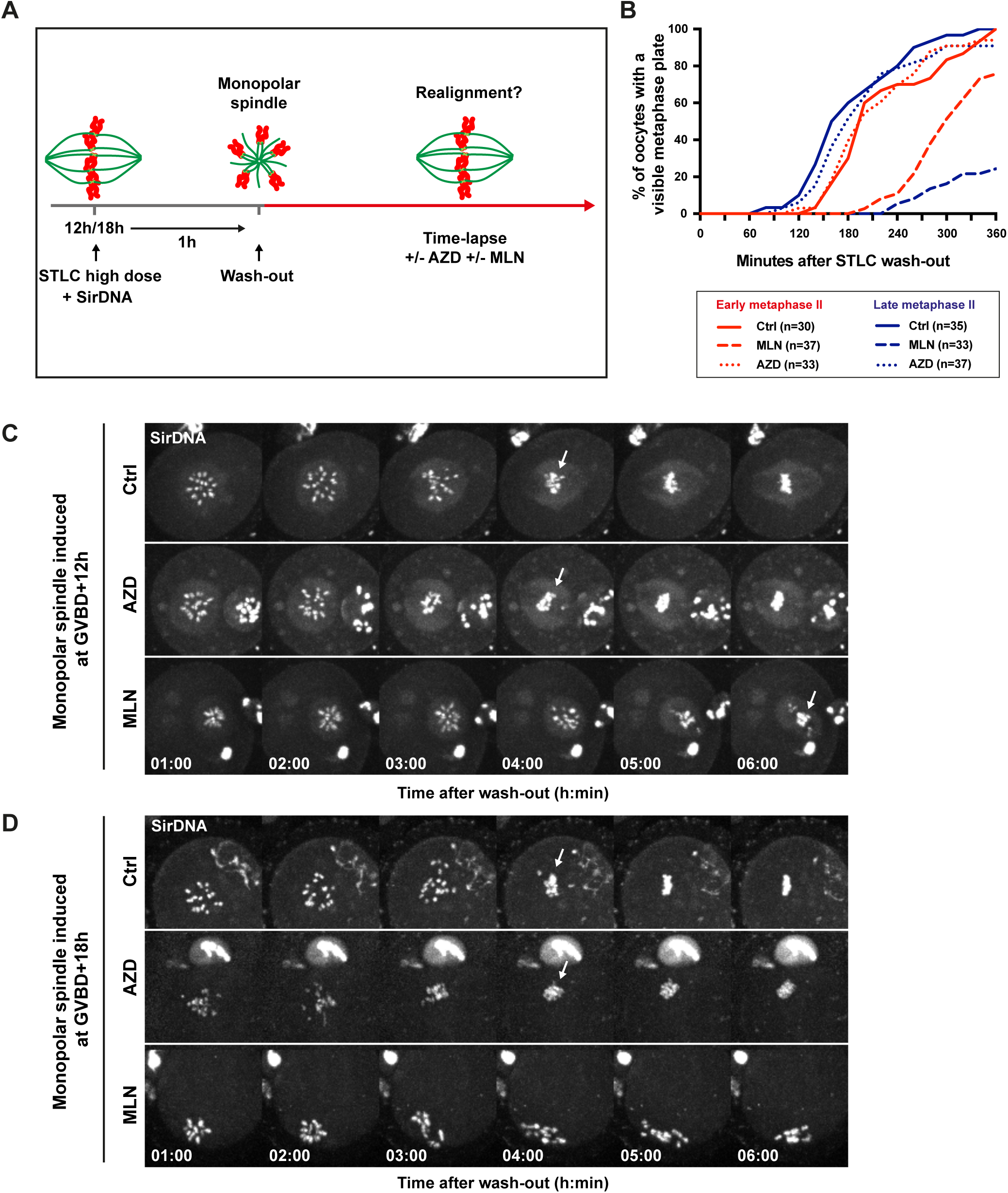
Monopolar attachments can be corrected in early and late metaphase II through Aurora A activity. (A) Scheme illustrating the experimental protocol of the experiment: oocytes were treated with elevated concentrations of STLC (4µM) at GVBD+12h or GVBD+18h to induce monopolar spindles. After 1 h, oocytes were washed to remove STLC, in the absence or presence of MLN8237 at 1 µM or AZD1152 at 0,1 µM. Chromosomes were stained with SirDNA throughout the procedure. The realignment of chromosomes was followed by time-lapse microscopy. (B) The time required for chromosomes to realign after wash-out was quantified for each oocyte by time-lapse microscopy. For each condition, the cumulative percentage of oocytes with a visible metaphase plate (some chromosomes still unaligned) relative to the time after wash-out is shown. n is the number of oocytes analyzed from 3 independent experiments. (C-D) Montage of representative movies obtained after induction of monopolar spindle at GVBD+12h or GVBD+18h and wash-out in the absence or presence of AZD1152 at 0,1 µM or MLN8237 at 1 µM. The time after wash-out is indicated (h:min). Arrows indicate when the metaphase plate appears.

In Figure 4B, we established at what time after release from STLC the metaphase plate became visible, even if some chromosomes were not aligned. Surprisingly though, after STLC removal, monopolar attached sister chromatids realigned even more efficiently in late than in early metaphase II. Indeed, 50% of the oocytes exhibited a visible metaphase plate after 180 min in early metaphase II, and after 150 min in late metaphase II (Figure 4B: solid curves, Figures 4C and 4D: first lines and Videos 10 and 11). This suggests that oocytes are able to convert monopolar attachments of sister chromatids into bipolar attachments even in late metaphase II. Thus, under these experimental conditions, the high levels of PP2A in late metaphase II are not sufficient to stabilize monopolar attachments.

How can this result be reconciled with the inefficient late metaphase II error correction described above? We hypothesized that Aurora A, which is located at spindle poles, contributes to the correction of attachment errors on monopolar spindles, even though Aurora A is not required for error correction on bipolar spindles, as we showed before. Indeed, it has been shown that Aurora A can phosphorylate the same kinetochore substrates as Aurora B/C on chromosomes close to the poles^39–41^. On a monopolar spindle, chromosomes are in close proximity to the pole^26^. To investigate the contribution of Aurora A in addition to Aurora B/C for the correction of monopolar attachments in our experiments, we released oocytes from STLC and allowed spindles to bipolarize in the presence of MLN8237 or AZD1152, inhibitors of Aurora A and Aurora B/C, respectively. Using 0.1 µM of AZD1152, a dose reported to inhibit only Aurora B/C but not Aurora A^36^, monopolar attached sister chromatids were mostly able to realign at the same time as untreated oocytes in early and late metaphase II (Figure 4B: dotted lines). Note that chromosome realignment was often not perfect compared to control oocytes, but a metaphase plate was always visible after Aurora B/C inhibition, even if a few chromosomes were still unaligned at the end of the movie (Figures 4C and 4D: second lines and Videos 10 and 11). However, when Aurora A was inhibited, sister chromatids that were attached in a monopolar manner completely failed to realign upon release from STLC, in both early and late metaphase II. In most cases, a metaphase plate did not form at all (Figures 4C and 4D: third lines and Videos 10 and 11), showing that Aurora A is contributing significantly to the correction of attachments on monopolar spindles. Crucially, when visible, the time of metaphase plate appearance was more affected in late than in early metaphase II. Indeed, 50% of early metaphase II oocytes compared to 20% of late metaphase II oocytes formed a metaphase plate within 5 h (Figure 4B: dashed lines). This result demonstrates that in the absence of Aurora A activity, Aurora B/C is able to counteract PP2A more efficiently in early than in late metaphase II, even on monopolar spindles.

In conclusion, our data show that the accumulation of PP2A in late metaphase II weakens the correction of faulty attachments by Aurora B/C on a bipolar spindle. On monopolar spindles however, the additional contribution of Aurora A allows correction of the most severe errors, even in late metaphase II.

### Low tension is also not detected in late metaphase II after *in vivo* maturation

Finally, we wanted to address whether the differences in error correction efficiency could be reproduced in metaphase II oocytes that have been matured *in vivo*, e.g., harvested after ovulation from the oviducts of female mice stimulated by injection of PMSG and hCG. Unfortunately, the exact timing of GVBD after hCG injection is unknown and may vary, but Thomas et al. recently estimated that GVBD and ovulation occur at 4 h and 12 h, respectively, after hCG addition in *ex vivo* culture of isolated follicles^42^. To have conditions comparable with our experiments performed after *in vitro* maturation, we collected *in vivo* matured oocytes from oviducts 14 h and 20 h after hCG injection.

When metaphase II oocytes were collected from the oviducts after *in vivo* maturation, unaligned chromosomes were only rarely observed and metaphase plate width was similar 14 h and 20 h after hCG injection. Also, metaphase plate width was slightly smaller than after *in vitro* maturation, probably reflecting a better efficiency of chromosome alignment *in vivo* (Figure S4A-C).

Addition of STLC induced a reduction in spindle length (Figures 5A and 5B) and inter-KT distances (Figures 5C and 5D), confirming the efficiency of STLC treatment at both time points after *in vivo* maturation. Of note, STLC did not affect the attachment and alignment of chromosomes (videos 12 and 13). Importantly, when *in vivo* matured oocytes were treated with 1 µM STLC 14 h after hCG injection, anaphase II was slightly delayed after strontium addition, compared to control oocytes, and a lower percentage of oocytes underwent anaphase II (80% instead of 100% in control oocytes). Crucially, when STLC was added 20 h after hCG injection at a final concentration of 1 µM, 100% of oocytes completed anaphase II with comparable efficiency to control oocytes, and they did so even at 2 µM of STLC (Figure 5E, S4D and S4E and videos 14 and 15). This result suggests that like *in vitro* matured oocytes, *in vivo* matured oocytes can detect and respond to low tension, but only at an early stage, not later. However, fewer oocytes were able to respond to low tension compared to *in vitro* matured oocytes, probably because oocytes are less synchronized after *in vivo* maturation. It was technically impossible to obtain *in vivo* matured oocytes earlier, as ovulation occurred approximately 12 hours after hCG injection.

**Figure 5.**
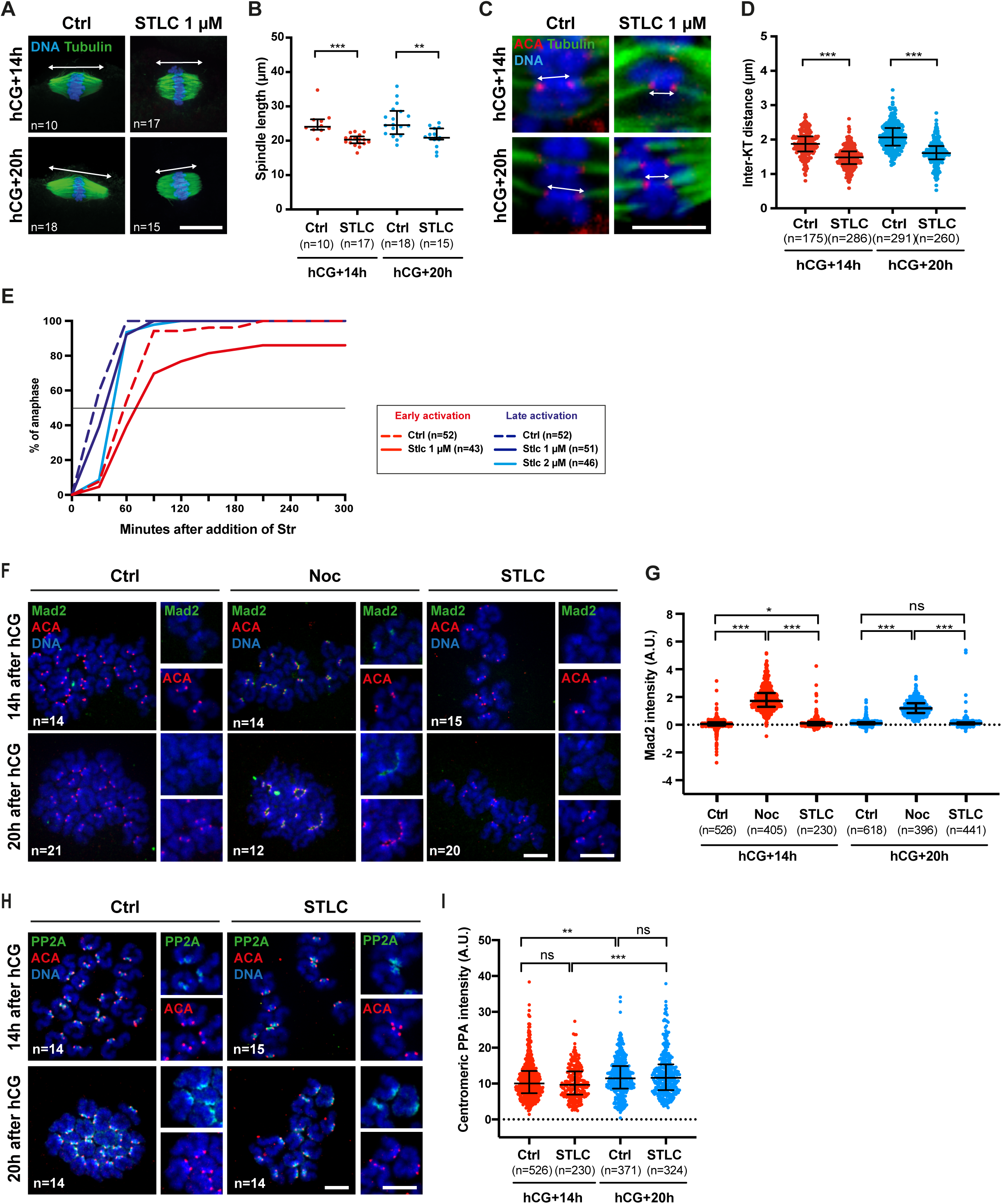
Low tension is also not detected in metaphase II after *in vivo* maturation. (A-D) *In vivo* matured meiosis II oocytes were collected in oviducts 14 h or 20 h after hCG injection and were left untreated (Ctrl) or treated with STLC 1 µM for 1 h, fixed and stained with DAPI (blue), anti-Tubulin antibody (green) and ACA (red). (A) The double-headed arrow indicates the spindle length. Scale bar: 20 µm. n: number of oocytes. (B) Graph showing the spindle length measured in each oocyte from (A). Median and interquartile range are indicated. n: number of oocytes from at least 3 independent experiments. (C) Magnifications of sister chromatid pairs from oocytes in (A). The double-headed arrow indicates the inter-kinetochore distance. Scale bar: 5 µm. (D) Graph showing inter-kinetochore (inter-KT) distances measured in each oocyte. Median and interquartile range are indicated. n: number of sister chromatid pairs analyzed in oocytes from (A) from at least 3 independent experiments. (E) *In vivo* matured meiosis II oocytes were collected in oviducts 14 h (red curves) or 20 h (blue curves) after hCG injection and incubated with SirDNA. Oocytes were left untreated (Ctrl) or treated with STLC (STLC) 1 or 2 µM, for 1h, before induction of anaphase II by strontium addition. The percentage of oocytes undergoing anaphase II relative to the time after addition of strontium was determined by time-lapse microscopy. The straight horizontal line represents 50% of the oocytes that have entered anaphase II. n: number of oocytes. (F) *In vivo* matured oocytes collected at 14 h or 20 h after hCG injection, were left untreated (Ctrl) or treated with Nocodazole 0,4 µM (Noc) or STLC 1 µM for 1 h. Chromosomes were spread and stained with ACA (red), anti-Mad2 antibody (green) and DAPI (blue). Smaller images correspond to magnifications of chromosomes. Scale bars: 10 µm. n: number of oocytes. (G) The mean intensity of Mad2 signal per centromere in oocytes from (F) has been quantified and normalized to the mean intensity of ACA (AU: arbitrary unit). The values from 3 independent experiments are shown and median and interquartile range are indicated. p values have been calculated by using Mann-Whitney’s test (*** p<0.0001, **p< 0.001, *p<0.05). n: number of centromeres. (H) *In vivo* matured oocytes collected at 14 h or 20 h after hCG injection, were left untreated (Ctrl) or treated with STLC 1 µM for 1 h. Chromosomes were spread and stained with ACA (red), anti-PP2A antibody (green) and DAPI (blue). Smaller images correspond to magnifications of chromosomes. Scale bars: 10 µm. n: number of oocytes. (I) The mean intensity of PP2A signal per centromere in oocytes from (H) has been quantified and normalized to the mean intensity of ACA (AU: arbitrary unit). The values from 3 independent experiments are shown and median and interquartile range are indicated. p values have been calculated by using Mann-Whitney’s test (*** p<0.0001, **p< 0.001, *p<0.05, ns: not significant). n: number of centromeres.

Mad2 was recruited to kinetochores after addition of nocodazole, confirming functionality of the SAC also in *in vivo* matured oocytes. However, no increase of Mad2 was detected after addition of STLC at 14 h or 20 h after hCG injection (Figures 5F and 5G), showing that *in vivo*, low tension is unable to induce efficient microtubule release and SAC activation even at an early stage. Alternatively, transient recruitment of Mad2 to kinetochores may not be detected at an early stage of *in vivo* maturation because the oocytes are not sufficiently synchronized.

As for *in vitro* matured oocytes, ineffective tension sensing was associated with high levels of PP2A that accumulate already at 14 h after hCG injection (Figures 5 H and 5I). The level of PP2A observed at 14 h after *in vivo* maturation was comparable to that observed at 18h after GVBD (10 A.U in Figure 3E and 7 A.U. in Figure 5I). This indicates that *in vivo* matured oocytes 14 h after hCG injection have already accumulated too much PP2A at kinetochores for efficient error correction. Thus, even though some *in vivo* matured oocytes show a response to low tension in early metaphase II, most oocytes do not, suggesting that error correction becomes inefficient earlier in these *in vivo* matured oocytes. Thus, these results show that it is not the *in vitro* maturation that weakens the error correction mechanism.

## DISCUSSION

Oocyte meiosis II faces specific challenges such as dealing with segregation errors carried over from meiosis I, or maintaining sister chromatid biorientation for an extended period of time during metaphase II arrest to await fertilization. During metaphase II arrest, chromosomes must remain aligned on the bipolar spindle for hours until fertilization. The time window during which fertilization can take place, before postovulatory aging and oocyte deterioration occur, is quite long (10-12 h post-ovulation in the mouse, and 24 h in human oocytes)^43,44^. Thus, the tension-sensing mechanism supposed to detect attachment errors, should be required in meiosis II to prevent post-fertilization aneuploidy. However, we confirmed here the persistence of unaligned chromosomes in metaphase II, as reported by others^28^, suggesting that incorrect attachments may not be detected and repaired efficiently.

Our data demonstrate that SAC functionality is well preserved throughout this prolonged metaphase II arrest, but that oocytes are only able to correct attachments during a limited temporal window. After *in vitro* maturation, low-tension attachments are well detected by the Aurora B/C-dependent error correction pathway in early metaphase II, but escape error correction in late metaphase II. After *in vivo* maturation, the window of active correction was more difficult to observe, suggesting that low-tension attachments are even better tolerated and do not result in SAC activation.

A recent study has argued that the SAC is able to block anaphase II onset after severe spindle alteration, such as produced by nocodazole, but cannot be activated by a small number of genetically induced unaligned chromosomes^45^. These observations are in agreement with our data in late metaphase II, corresponding to the cell cycle stage examined in the study ^45^. First, SAC response is only transient, and oocytes will eventually continue cell cycle progression even in the presence of wrongly attached kinetochores. Once in late metaphase II, failure to activate the SAC can be explained by a defective tension-sensing mechanism, as we propose here.

We show that oocytes turn off error correction in late metaphase II by shifting the balance between PP2A and Aurora B/C in favor of PP2A. PP2A-B56 recruitment to kinetochores is mediated by the SAC components BubR1 and Mps1, and we have previously shown that at least Mps1 remains associated with kinetochores in anaphase II^37,46^, supporting that PP2A is continuously recruited to kinetochores throughout metaphase II arrest. PP2A has been also proposed to be required to protect cohesin until anaphase II onset, and to be inhibited in anaphase II by Set/I2PP2A to deprotect cohesin^47^. However, our recent unpublished data show that in mouse oocytes, cohesin protection is absent during the prolonged metaphase II arrest, and that the role of Set/I2PP2A is independent of PP2A. Future work will determine which pool of PP2A counteracts error correction in metaphase II, but we hypothesize that the PP2A pool at centromeres ensures microtubule stabilization via Hec1 dephosphorylation, whereas the PP2A pool between sister chromatids may be required for other functions.

Our results highlight a major difference between meiosis I and meiosis II, since low tension allows Aurora B/C to counteract PP2A in metaphase I^26^, but not in late metaphase II. This could be explained by differences in PP2A levels, which are much higher in metaphase II than in metaphase I^27^. Accordingly, PP2A is found around centromeres and between sister centromeres in metaphase II, but is more restricted between sister centromeres in metaphase I^37,47^. We propose that in late metaphase II, low tension is better tolerated because of the accumulation of more PP2A at centromeres counteracting Aurora B/C. As a trade-off, errors that are not repaired early in metaphase II, persist until late metaphase II.

Unexpectedly, after induction of monopolar spindles, late metaphase II oocytes converted monopolar into bipolar attachments very efficiently, thus demonstrating that in principle, low tension attachments can be corrected. However, we show here that it is mainly Aurora A that is required to restore chromosome biorientation after induction of monopolar spindles, in both early and late metaphase II oocytes. Thus, on monopolar spindles, attachments can be removed by Aurora A-dependent destabilization activity. Indeed, Aurora A has been shown to locally phosphorylate Hec1 on kinetochores of chromosomes that are found near the spindle poles, contributing to destabilize incorrect attachments^39–41^. In contrast, Aurora A is not contributing to error correction on bipolar spindles in metaphase II. The combined activity of Aurora A at poles and Aurora B/C at kinetochores is probably more efficient to counteract PP2A on monopolar spindles, where chromosomes are found closer to the poles. However, when Aurora A is inhibited, monopolar attachments are less well repaired in late than in early metaphase II, further confirming that Aurora B/C is much more efficient in early metaphase II.

When comparing *in vivo* with *in vitro* matured oocytes, we were not able to define exactly the time window during which correction is still possible *in vivo*. It seems that this window is already overwhelmed once the oocytes are ovulated. Thus, *in vivo*, oocytes with ongoing error correction could not be fertilized because they would not have reached the oviduct, whereas *in vitro*, oocytes can enter anaphase II as early as 12h after GVBD. It remains to be determined whether hormonal stimulation accelerates progression of oocyte through metaphase II, which would not help to define the window of active error correction *in vivo*.

Overall, our data using mouse oocytes show a higher susceptibility of late metaphase II oocytes to maintain incorrectly attached chromosomes when efficient fertilization is still possible *in vivo* and *in vitro*. The physiological reason why error correction in oocytes weakens in metaphase II is unclear. It is possible that PP2A needs to accumulate in metaphase II for other functions and indirectly affects error correction. It is also possible that PP2A accumulation is required to stabilize microtubule attachments over a long period of time. Indeed, PP2A accumulation has been shown to be required to stabilize microtubule attachments in metaphase of meiosis I, by counteracting Aurora B/C, which remains in close proximity to kinetochores^27^. Without PP2A accumulation, Aurora B/C may continue to destabilize microtubule interaction even when they are correctly attached. Thus, ongoing error correction during metaphase II arrest would be more deleterious and lead to more errors. In contrast, transient activation followed by a weakening of error correction would allow only a few incorrect attachments to escape and be stabilized by PP2A. The stabilization of microtubule attachments by PP2A is likely to be very critical in metaphase II, when chromosomes must remain attached for a long time while awaiting fertilization. Finally, weakening error correction during metaphase II should protect rather than induce aneuploidy, at least in young mice.

Importantly, in mouse and human oocytes, segregation errors in meiosis II increase with maternal age, due to an increase in misaligned chromosomes, the prevalence of single chromatids, or loss of kinetochore integrity^48–52^. Thus, turning off error correction during metaphase II arrest would have more severe consequences in oocytes from aged mice or older women, since there would be more errors to deal with, and likely more errors persisting until late metaphase II. Importantly, oocytes from aged mice fail to correct aberrant attachments that are under less tension in metaphase II, supporting the inefficiency of error correction^45,51^. Thus, error correction seems to be functional sufficiently long for correct segregation in oocytes that are not facing additional challenges, such as increased maternal age for example. Insights into the functionality of error correction relative to the time spent in metaphase II arrest as well as maternal age may contribute to better determine the optimal temporal window for fertilization also in human oocytes, to minimize aneuploidies and at the same time, achieve optimal fertilization rates^53^.

## ACKNOWLEDGEMENTS

We are grateful to Sandra Touati and Aude Dupré for discussions and comments on the manuscript. We thank Anastasia Shihabi and Maya Merabet for their help during their undergraduate internships, and members of the MOM group for advice and discussion. We thank Nicolas Minc for the use of the micropipette puller. We acknowledge the ImagoSeine core facility of the Institut Jacques Monod, member of the France BioImaging infrastructure (ANR-10-INBS- 04) and GIS-IBiSA. We thank the members of the animal house and the informatics and administrative support services of the Institut Jacques Monod.

AL received a 3-year PhD fellowship from Sorbonne Université (ED393) and a 1-year PhD fellowship from the Labex “Who Am I” (ANR-11-LABX-0071; Idex ANR-18-IDEX-0001). The KW lab was funded by the Agence Nationale de la Recherche (ANR-19-CE13-0015, ANR-23-CE13-0015- 01), La Fondation de la Recherche Médicale (Equipe FRM DEQ 202103012574) and core funding from the CNRS, Université Paris Cité and Sorbonne Université.

## AUTHOR CONTRIBUTIONS

Experiments were performed by A.L, A.K-S and E.B. Figures were prepared by E.B. The manuscript was written by E.B. with help from K.W. and input from all authors. Funding was obtained by K.W, supervision and project administration were done by E.B. and K.W.

## STAR METHODS

### RESOURCE AVAILABILITY

#### Lead contact

Further information and requests for resources and reagents should be directed to and will be fulfilled by the lead contact, Katja Wassmann.

#### Materials availability

All unique reagents generated in this study are available without restriction from the lead contact. No new mouse lines or plasmids were generated in this study.

#### Data and code availability

Microscopy data reported in this paper will be shared by the lead contact upon request. This paper does not report original code.

Any additional information required to reanalyze the data reported in this paper is available from the lead contact upon request.

### EXPERIMENTAL MODEL AND STUDY PARTICIPANT DETAILS

#### Animals

Female CD-1 mice aged 7-9 weeks were purchased (Janvier, France) and maintained with *ad libitum* access to food and water in the conventional mouse facility of UMR7622 (authorization C75-05-13), IBPS (authorization A75-05-24) and UMR7592 (authorization C75-13-17). The project was approved by ethical review under French law (authorization APAFIS#50208- 202406051134677v3). To collect prophase I oocytes from ovaries, mice were not exposed to any drug treatment or procedure before being sacrificed by cervical dislocation between 8 and 16 weeks of age. To collect *in vivo* matured oocytes, mice were injected with 5 units of PMSG (Pregnant Mare Serum Gonadotropin, Abbexa LTD, abx260389), and 48 h later with 5 units of hCG (human Chorionic Gonadotropin, Abbexa LTD, abx260092) to stimulate *in vivo* maturation and ovulation. 14 h or 20 h after hCG injection, mice were sacrificed by cervical dislocation, and oocytes were collected in the oviducts.

## METHOD DETAILS

### Mouse oocyte culture

Prophase I arrested oocytes, visualized by the presence of a germinal vesicle (GV), were collected from ovaries after dissection in homemade M2 medium ^54^. M2 medium was supplemented with 100 µg/mL dibutyryl cyclic AMP (dbcAMP) (Sigma-Aldrich, D0260) to maintain oocytes in prophase I arrest. Where indicated, prophase I oocytes were injected with specific mRNAs (see below). Oocytes were released from prophase I arrest after several washes in M2 medium without dbcAMP. Germinal vesicle breakdown (GVBD) was determined by the disappearance of the GV, which is easily visible under binocular. Oocytes undergoing GVBD within 90 minutes were used for the experiments. After GVBD, *in vitro* maturation was performed in drops of M2 medium covered with mineral oil (Fujifilm Irvine Scientific, 9305) and incubated at 37°C without CO2, or in drops of M16 medium (MERK Millipore, MR-10P) covered with mineral oil and incubated at 37°C with 5% of CO2. M16 is necessary when metaphase II arrested oocytes need to be activated with strontium chloride to induce anaphase II without fertilization. In this case, oocytes are incubated in CaCl_2_-free M2 medium for 30-60 min and transferred to CaCl_2_-free M2 containing 10 mM of strontium chloride (Sigma-Aldrich, 204463).

*In vivo* matured oocytes were collected from the oviducts, 14 h or 20 h after hCG injection: the cumulus containing oocytes were released from the ampulla in M2 medium, and dissociated in M2 containing 0,125 µg/µl of hyaluronidase (Sigma-Aldrich, H4272). After 5 min, individualized metaphase II oocytes were collected and washed several times in M2 medium.

### Drug treatment

The following drugs were dissolved in DMSO (Sigma-Aldrich, D2650) and were used at the indicated final concentrations in M2 medium: STLC 1 µM, 2 µM or 4 µM (S-Trityl-L-cystein, Sigma-Aldrich, 164739), Nocodazole 0.4 µM (Sigma-Aldrich, M1404), AZD1152-HQPA 0.1 µM or 0.5 µM (Sigma-Aldrich, SML0268), Reversine 0.5 µM (Interchim, 10004412), MLN8237 1 µM (Clinisciences, A4110). Reversine is also added to the mineral oil covering the drops to compensate for drug diffusion into the oil, and oocytes have to be treated and cultured in separate dishes.

### Fixation of whole oocytes and chromosome spreads

The zona pellucida surrounding the oocyte was dissolved in successive drops of Tyrode’s acid solution (NaCl 137 mM, KCl 2.7 mM, CaCl2 1.8 mM, MgCl2 0.5 mM, D-Glucose 5.5 mM, PVP 0.09 mM, pH 2.3). Fixation of whole oocytes was performed as previously described ^26,54^. In short, oocytes were transferred to glass bottom chambers coated with concanavalin A (Sigma-Aldrich) in M2 medium containing 0.2% PVP (Polyvinylpyrrolidone, Merk Millipore) before slow centrifugation at 1400 rpm for 13 min at 37°C. For spindle staining, oocytes were subjected to cold treatment for 6 min in 80 mM PIPES, 1 mM MgCl2, pH7.4 in a dish placed on ice. Oocytes were fixed for 30 min in 1X BRB80 (80 mM PIPES, 1 mM MgCl2, 1 mM EGTA, pH7.4) containing 0.3% Triton-100X and 1.9% formaldehyde (Sigma-Aldrich F1635). After washes with 1X PBS, oocytes were permeabilized and saturated overnight at 4°C in 1X PBS, 3% BSA, 0.1% Triton-X100. For chromosome spreads, oocytes were fixed in a hypotonic solution (H20, 1% paraformaldehyde (Sigma-Aldrich 441244), 0.15% Triton-X100, 3 mM DTT), which allows the membrane to dissolve and to spread the chromosomes on a 10-well slide (Fischer Scientific-10588681).

### Immunostaining

After saturation with 1X PBS, 3% BSA, fixed whole oocytes or chromosome spreads were incubated with primary antibodies for 2-3 h, washed and incubated with secondary antibodies for 1h30, at room temperature. The following antibodies were used at the indicated concentrations: Human anti-centromere protein antibody (ACA, Clinisciences, 15-234-0001, 1:100), mouse monoclonal anti-αTubulin conjugated with FITC (DM1A, Sigma-Aldrich, F2168, 1:100), rabbit polyclonal anti-Hec1pS55 (Euromedex, GTX70017, 1:100), rabbit polyclonal anti-Mad2 (1:100) ^55^, mouse monoclonal anti-PP2A C-subunit clone 1D6 conjugated with Alexa Fluor 488 (Sigma-Aldrich, 05-421-AF488, 1:100), rabbit polyclonal anti-Aurora C (Thermofisher, PA5118973, 1:100). Donkey anti-human Cy3 (Jackson ImmunoResearch, 709-166-149, 1:200), Donkey anti-rabbit AF488 (Jackson ImmunoResearch, 711-546-152, 1:200), Donkey anti-mouse AF488 (Jackson ImmunoResearch, 715-546-150, 1:200), Donkey anti-Rabbit AF647 (Jackson ImmunoResearch, 711-606-152). Slides were mounted using VECTASHIELD mounting medium supplemented with DAPI (Eurobio H-1200).

### *In vitro* transcription and oocyte microinjection

H2B-RFP^56^, Securin-YFP^57^ or Tubulin-GFP^58^ coding sequences inserted into the pRN3 plasmid were used for *in vitro* transcription. Plasmids were digested with SfiI restriction enzyme, and the resulting linearized plasmid was purified by phenol/chloroform purification. *In vitro* transcription was performed by using the mMessage mMachine T3 Kit (Invitrogen, AM1348). After 2 h of transcription, mRNAs were purified by using the RNeasy Mini Kit (Qiagen, 74104).

Purified mRNAs were injected into prophase I arrested oocytes using a FemtoJet microinjector pump (Eppendorf) with continuous flow, on an inverted Nikon Eclipse Ti microscope. Microinjection pipettes were made by pulling capillaries (Harvard Apparatus, 30-0038) with a magnetic puller (Narishige, PN-31 or Sutter Instrument, P-1000) and oocytes were manipulated using a holding pipette (BIOZOL, Eppendorf, 5178108.000).

### Image acquisition and live imaging

Chromosome spreads and fixed whole oocytes were observed using a 100X/1.4 NA oil objective, on a Zeiss Axiovert 200M inverted microscope, or a Nikon Eclipse Ti2-E inverted microscope, both coupled to an EMCCD camera (Evolve 512, Photometrics) and a Yokogawa CSU-X1 spinning disc. 6-10 Z-sections spaced by 0.4 µm were acquired for chromosome spreads and 25-30 Z-sections spaced by 0.4 µm were acquired for whole oocytes. Wavelengths of 491 nm, 561 nm, 642 nm and 405 nm were used for detection of AlexaFluor488, Cy3, AlexaFluor647, and Dapi, respectively.

Live imaging from Figure 1B (Video 1) were performed with a 63X/1.4NA oil objective, on a Zeiss Axiovert 200M inverted microscope, coupled to an EMCCD camera (Evolve 512, Photometrics) and a Yokogawa CSU-X1 spinning disc. Wavelengths of 491 nm and 561 nm were used for detection of Tubulin-GFP and H2B-RFP, respectively.

Live imaging from Figures S1, S3 and S4 (Videos 2, 3, 5, 6, 7, 9, 14 and 15) was performed with a Plan APO 20X/0.75 NA objective, on a Nikon Eclipse Ti2-E inverted microscope coupled to a Prime sCMOS camera (Photometrics) and a PrecisExite High PowerLED fluorescence. RFP, YFP or Far-red channels were used for detection of H2B-RFP, Securin-YFP and SirDNA, respectively.

Live imaging from Figure 4 (Videos 10, 11) was performed with a Plan APO 20X/0.75 NA objective, on a Nikon Eclipse Ti2-E inverted microscope, coupled to an EMCCD camera (Evolve 512, Photometrics) and a Yokogawa CSU-X1 spinning disc. Wavelength of 642 nm was used for detection of SirDNA.

All movies were performed under temperature-controlled conditions at 37°C. 15-20 Z-sections spaced by 3 µm were acquired every 10 min. Where indicated, chromosomes are visualized with SirDNA 1 µM (far-red DNA labelling probe, TEBU BIO, 251SC007) added to the culture medium 1 h prior to imaging. All Images were acquired with Metamorph software using the same setting for all conditions and replicated experiments.

Super-resolution images of fixed whole oocytes (Videos 4, 8, 12, 13) were acquired with an Elyra 7 SIM microscope from the Imagoseine facility, using a 63x/1.4 Plan-Apochromat oil objective DIC M27 (420762-9900-799), on a Zeiss Axio Observer 7 stand with definite focus 3, equipped with a 130 mm x 100 mm motorized piezo stage and a 500 µm piezo Z-stage, and coupled to two Edge 4.2 CLHS (PCO) cameras, set for simultaneous dual-channel display. 100-200 z-sections were acquired at 0.1 µm intervals. Wavelengths of 405 nm, 488 nm, and 561 nm were used for detection of DAPI, AlexaFluor488, and CY3, respectively. Super-resolution 3D Lattice SIM images were acquired and processed using SW ZEN Black 3.4 software.

Movies, composites and image projections were generated on Fiji software (NIH). Brightness and contrast were adjusted equally to all conditions compared.

## QUANTIFICATION AND STATISTICAL ANALYSIS

All quantifications were performed by using Fiji software.

### Signal quantification on chromosome spread

To quantify centromeric signals of Mad2, pS55Hec1 and PP2A (Fig2D, 3B, 3E, 5G), a circle of 10×10 pixels was placed around each centromere. Background signal was measured in a second circle placed on the chromosome adjacent to the centromeres. Mean fluorescence intensities were measured on SUM projections for channels corresponding to the protein of interest and to ACA. After background subtraction, mean fluorescence intensities were normalized to ACA signals from the same centromere. To quantify total PP2A and Aurora C signals (Fig3D, 3G), mean fluorescence intensities for the protein of interest and for ACA, were measured in manually selected areas including centromeres and between centromeres. Background signal was measured in the same areas placed on the chromosome. After background subtraction, mean fluorescence intensities of the protein of interest were normalized to ACA signals from the same area. Fluorescence intensities were plotted on a graph using GraphPad Prism 6 software, and statistical analysis was performed using the nonparametric Mann-Whitney U-test (ns: not significant, * p< 0.05, ** p< 0.001, *** p< 0.0001) and included at least three independent experiments.

### Signal quantification on time-lapse

To quantify Securin-YFP in time-lapse acquisitions (Fig2B, 2H), mean fluorescence intensities were measured in a square of 100×100 pixels placed inside the oocyte. Background signal was measured in a second square placed outside the oocyte. Mean intensities were measured on the MAX projection for each time point of the time-lapse. After background subtraction, the mean fluorescence intensities of Securin-YFP were normalized to the value obtained at t_0_ of the time-lapse. The mean values for each time point from at least three independent experiments were used to generate a kinetic curve using GraphPad Prism 6.

### Quantification of unaligned chromosomes

In Figures 1F, the number of DAPI-stained chromosomes clearly outside the metaphase plate was manually counted in each oocyte, from at least three independent experiments. A histogram showing the percentage of oocytes containing 0, 1-2, or more than 2 unaligned chromosomes was generated using GraphPad Prism 6 software.

### Quantification of distances

To quantify the metaphase plate width (Fig1E), a rectangle was manually defined on the MAX projection, closely around the centrally grouped chromosomes, excluding the unaligned chromosomes. The width of the rectangle was measured in each oocyte using Fiji software. To quantify spindle length (Figure S2B, S2G and 5B), a line was manually defined on the MAX projection, between the two spindle poles visualized by Tubulin staining. The line length was measured in each oocyte using Fiji software. To quantify inter-kinetochore distance (Figure S2D, S2H and 5D), a line was manually defined between two sister kinetochores stained with ACA. For better visibility of each sister kinetochore, inter-kinetochore distances were measured by considering each Z-section individually. Values obtained from at least three independent experiments were plotted on graphs using GraphPad Prism 6 software. Statistical analyses were performed using the nonparametric Mann-Whitney U-test (ns: not significant, * p< 0.05, ** p< 0.001, *** p< 0.0001).

## SUPPLEMENTAL FIGURES

**Figure S1.**
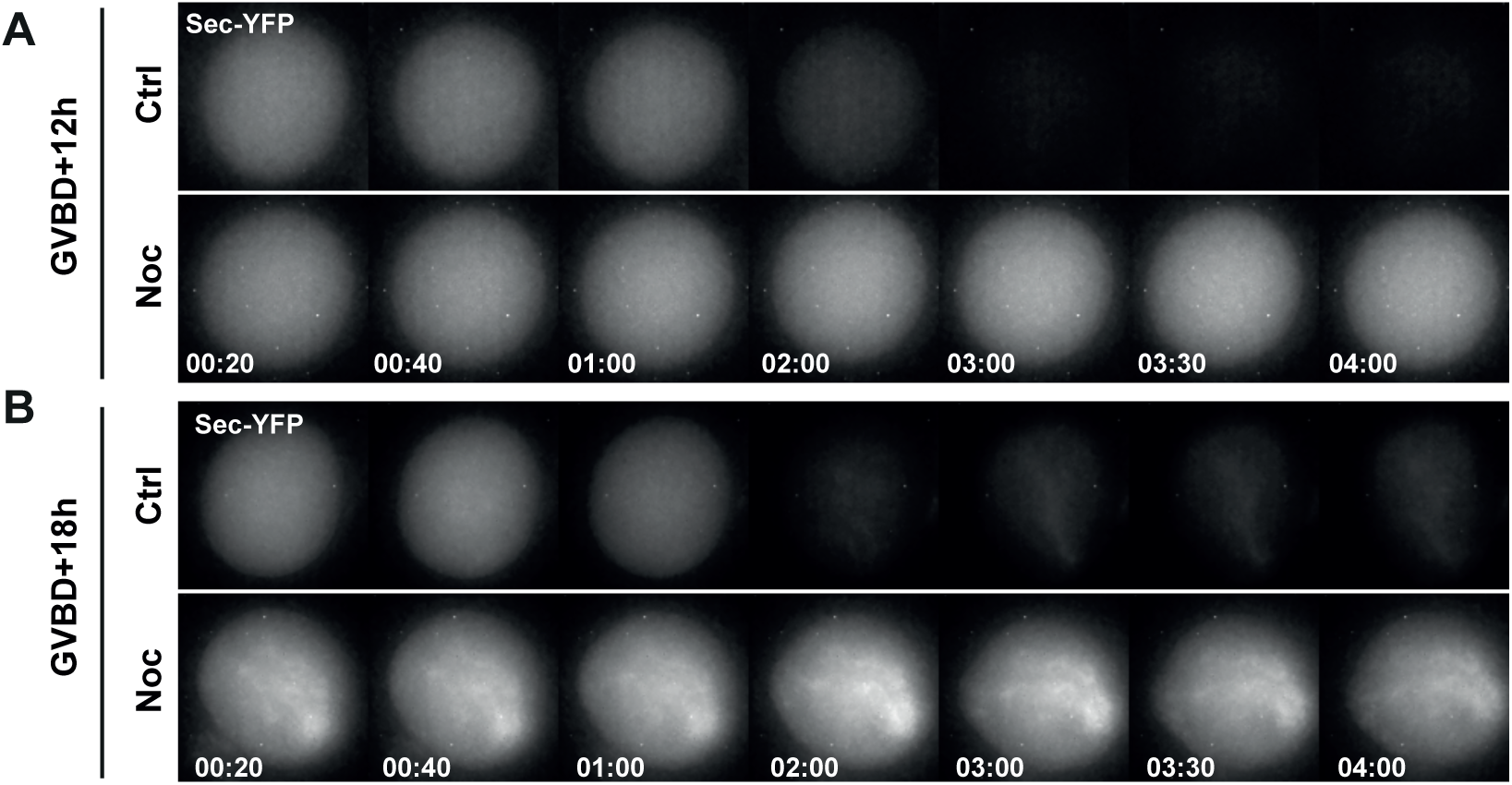
Anaphase II onset is inhibited by nocodazole when added in early or late metaphase. Montage of representative movies analyzed in Figure 2B. The time after addition of strontium is indicated (h: min). (A) Oocytes injected with Securin-YFP mRNA were left untreated (Ctrl), or treated at GVBD+12h with nocodazole 0,4 µM (Noc) for 1 h, and activated by addition of strontium. (B) Oocytes injected with Securin-YFP mRNA were left untreated (Ctrl) or treated at GVBD+18h with nocodazole 0,4 µM (Noc) for 1 h and activated by addition of strontium. (Related to Figure 2B)

**Figure S2.**
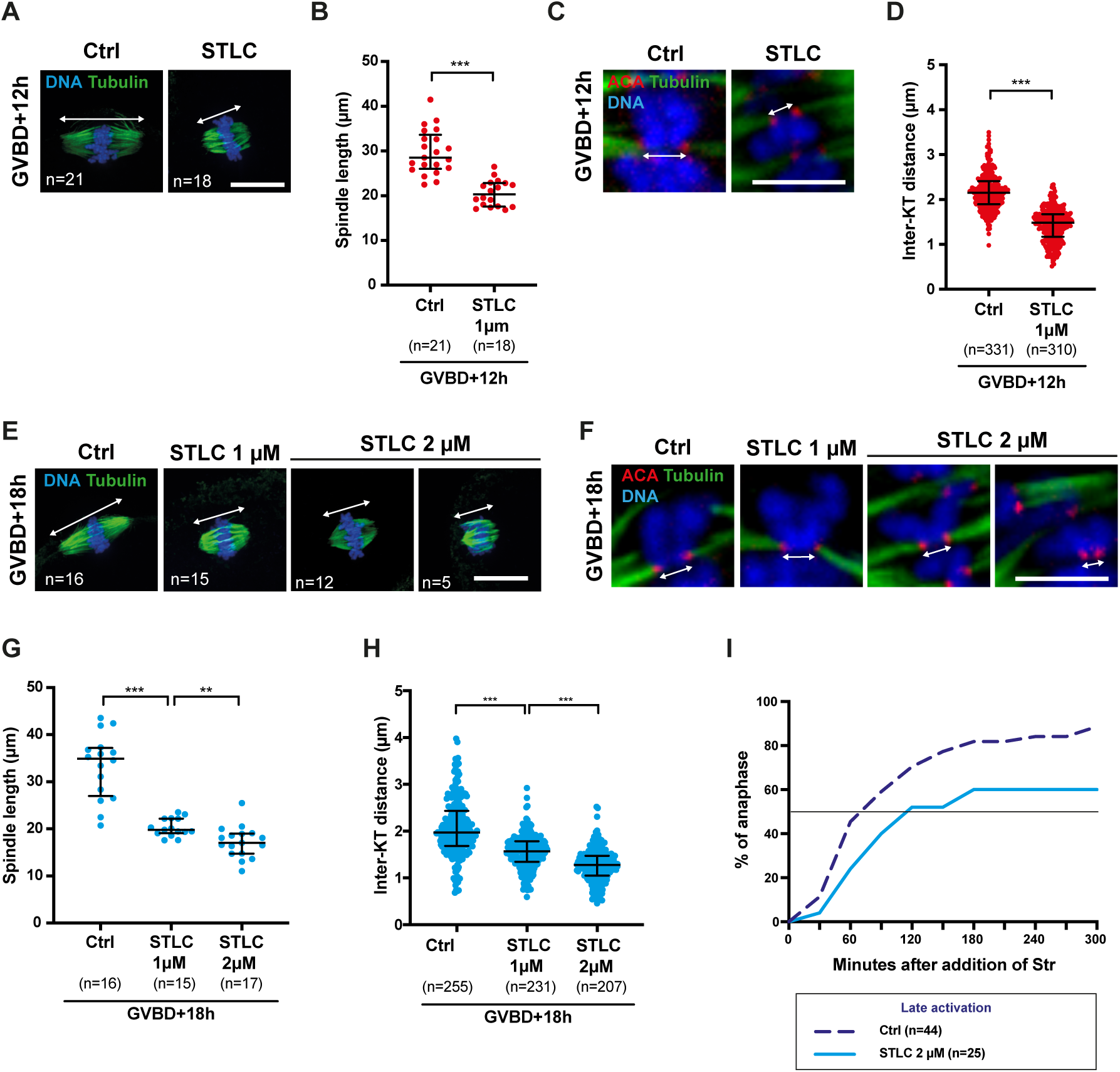
Inhibiting Eg5 induces short spindle and low tension in meiosis II. (A-D) *In vitro* matured meiosis II oocytes at 12h after GVBD (early metaphase II) were left untreated (Ctrl) or treated with STLC 1 µM for 1 h, fixed and stained with DAPI (blue), anti-Tubulin antibody (green) and ACA (red). (A) The double-headed arrow indicates the spindle length. Scale bar: 20 µm. (B) Graph showing the spindle length measured in each oocyte from (A). n: number of oocytes. (C) Magnifications of sister chromatid pairs of oocytes from (A). The double-headed arrow indicates the inter-kinetochore distance. Scale bar: 5 µm. (D) Graph showing inter-kinetochore (inter-KT) distances measured in each oocyte from (A). n: number of sister chromatid pairs analyzed in oocytes from (A) from 4 independent experiments for each condition. (E-H) *In vitro* matured meiosis II oocytes at 18h after GVBD (late metaphase II) were left untreated (Ctrl) or treated with STLC 1 or 2 µM for 1 h, fixed and stained with DAPI (blue), anti-Tubulin antibody (green) and ACA (red). (E) The double-headed arrow indicates the spindle length. At 2 µM STLC, a small fraction of oocytes undergoes spindle collapse. Scale bar: 20 µm. (F) Magnifications of sister chromatid pairs of oocytes from (E). The double headed-arrow indicates the inter-kinetochore distance. At 2 µM STLC, chromosomes may lose attachment as the spindle collapses. Scale bar: 5 µm. (G) Graph showing the spindle length measured in each oocyte from (E). n: number of oocytes. (H) Graph showing inter-kinetochore (inter-KT) distances measured in each oocyte from (E). n: number of sister chromatid pairs analyzed in oocytes from (E) from 3 independent experiments for each condition. In all the graphs, median and interquartile range are indicated. p values have been calculated by using Mann-Whitney’s test (*** p<0.0001, **p< 0.001, *p<0.05, ns: not significant). (I) *In vitro* matured oocytes incubated with SirDNA were left untreated (Ctrl) or treated with STLC 2 µM (STLC), for 1 h at GVBD+18 h. The percentage of oocytes undergoing anaphase II relative to the time after addition of strontium was determined by time-lapse microscopy. The straight horizontal line represents 50% of the oocytes that have entered anaphase II. n: number of oocytes. (Related to Figure 2)

**Figure S3.**
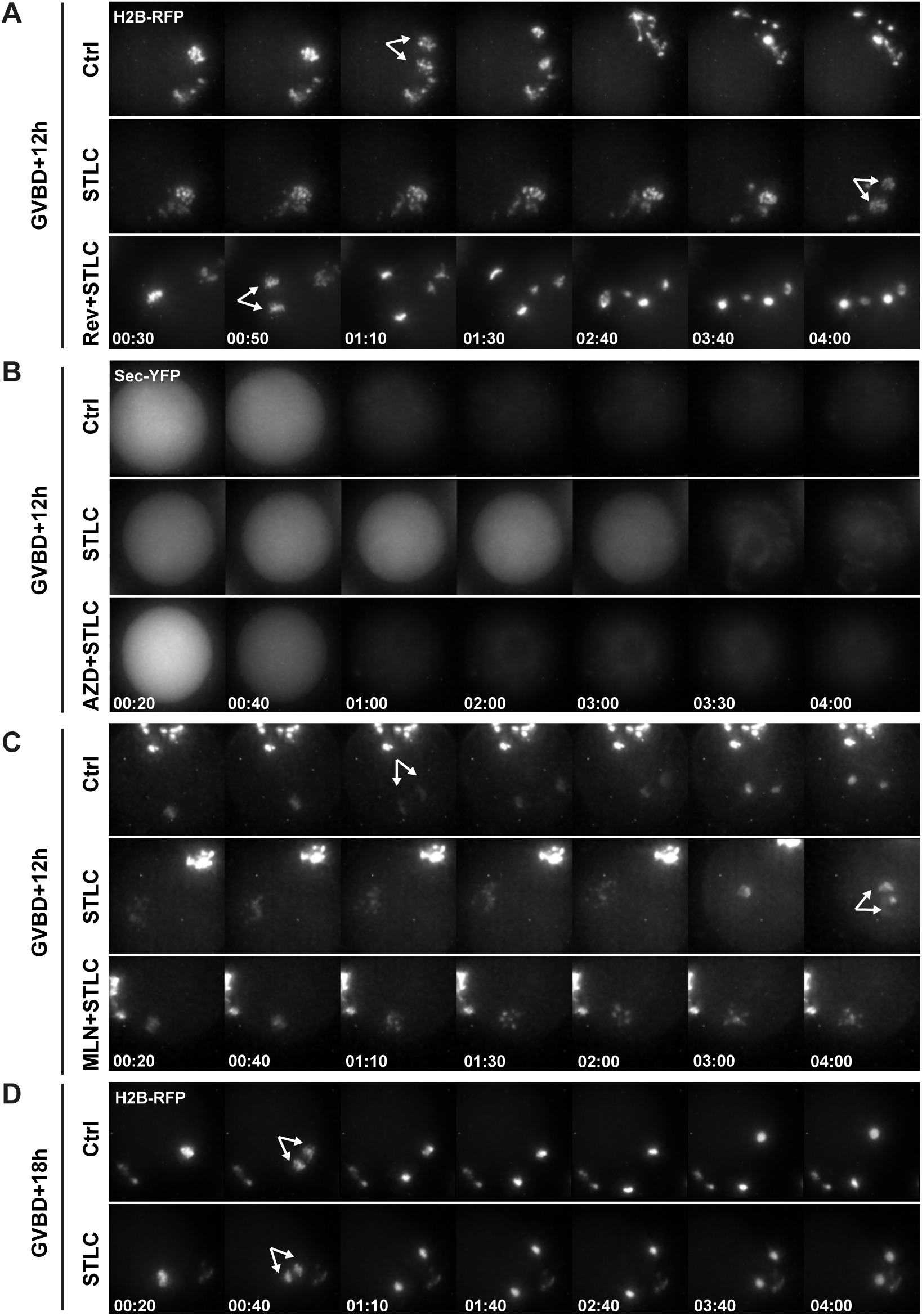
Anaphase II onset is delayed by STLC only when added in early metaphase II. Montage of representative movies analyzed in Figure 2 (E-H). The time after addition of strontium is indicated (h: min). Arrows indicate sister chromatid segregation and thus, anaphase II onset. (A) Oocytes injected with H2B-RFP mRNA were left untreated (Ctrl) or treated at GVBD+12h with STLC 1 µM, +/- Reversine 0,5 µM (Rev) for 1 h, and activated by addition of strontium. (B) Oocytes injected with Securin-YFP mRNA were left untreated (Ctrl), or treated at GVBD+12h with STLC 1 µM +/- AZD1152 0,5 µM (AZD) for 1 h, and activated by addition of strontium. (C) Oocytes incubated with SirDNA were left untreated (Ctrl) or treated at GVBD+12h with STLC 1 µM, +/- MLN8237 1 µM (MLN) for 1h, and activated by addition of strontium. (D) Oocytes injected with H2B-RFP mRNA were left untreated (Ctrl) or treated at GVBD+18h with STLC 1 µM for 1 h, and activated by addition of strontium. (Related to Figure 2 E-H)

**Figure S4.**
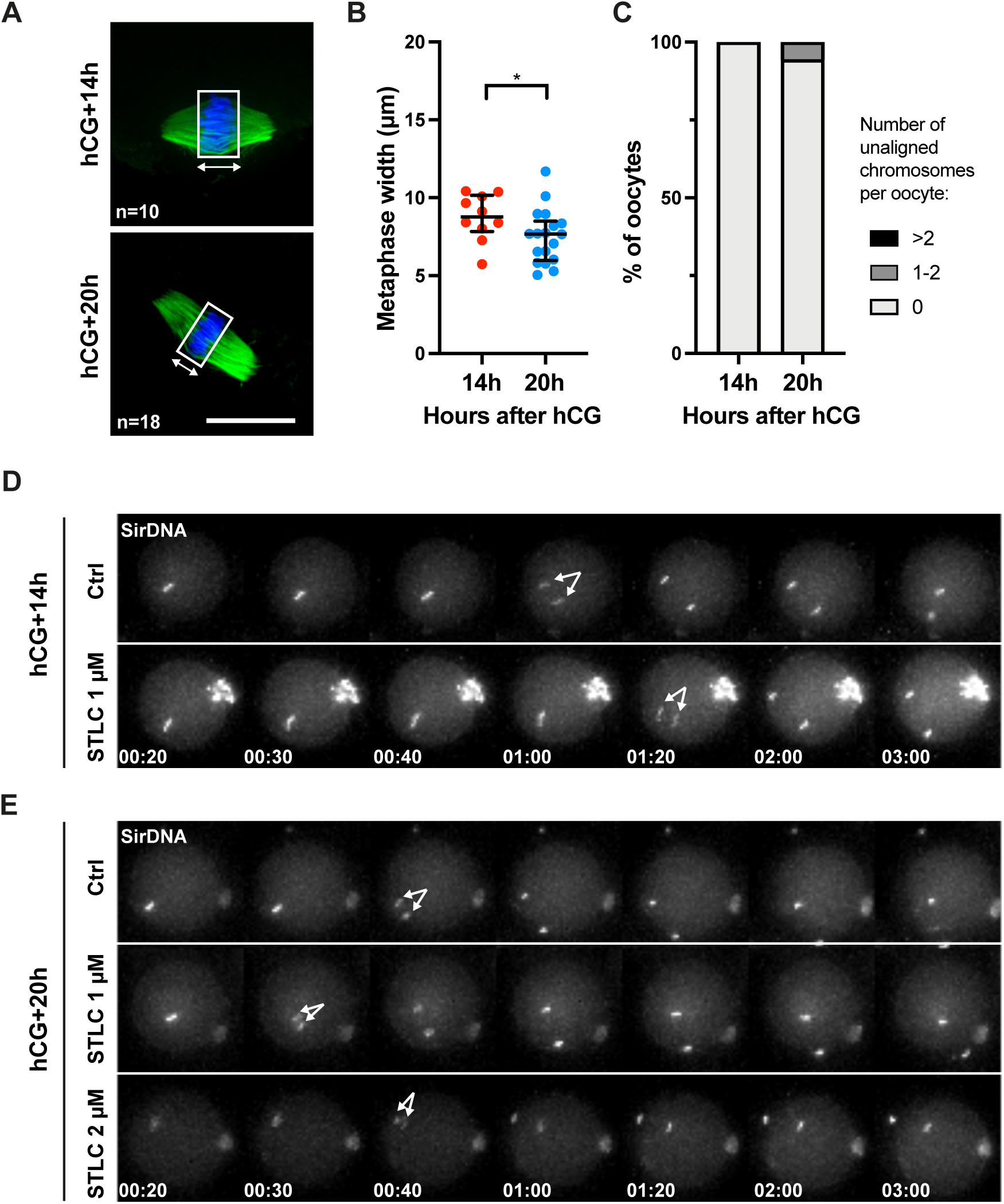
Anaphase II onset is not delayed by STLC after *in vivo* maturation. (A) *In vivo* matured oocytes were fixed in meiosis II at 14 h or 20 h after hCG injection and stained with DAPI (blue) and anti-Tubulin antibody (green). The rectangle defines the metaphase plate, and the double-headed arrow indicates how the width of the metaphase plate was measured in (B). Scale bar: 20 µm. n is the number of oocytes analyzed from at least 3 independent experiments for each condition. (B) The width of the metaphase plate was measured in each oocyte from (A). Median and interquartile range are indicated. p values have been calculated by using Mann-Whitney’s test (*p<0.05, ns: not significant). (C) The percentage of oocytes containing 0, 1-2, or more than 2 unaligned chromosomes was quantified in each oocyte from (A). Note that more than 2 unaligned chromosomes were never observed under our conditions. Chromosomes were considered unaligned if they were clearly outside the metaphase plate. (D-E) Montage of representative movies analyzed in Figure 5E. The time after addition of strontium is indicated (h: min). Arrows indicate sister chromatid segregation and thus, anaphase II onset. (D) *In vivo* matured oocytes collected at 14 h after hCG injection were incubated with SirDNA and left untreated (Ctrl) or treated with STLC 1 µM for 1 h, and activated by addition of strontium. (E) *In vivo* matured oocytes collected at 20 h after hCG injection were incubated with SirDNA and left untreated (Ctrl) or treated with STLC 1 or 2 µM for 1 h, and activated by addition of strontium. (Related to Figure 5)

## SUPPLEMENTAL MOVIES

**Video 1: Progression from metaphase I to metaphase II in control oocyte**

Example of an oocyte injected with H2B-RFP and Tubulin-GFP mRNA in prophase I. Acquisitions of RFP and GFP channels were started at GVBD+6h (h:min) every 15 min. Shown is the overlay from stack of 15 z-sections spaced by 3 µm. This is a representative movie from 23 oocytes from 3 independent experiments. Related to Figure 1B.

**Video 2: Visualization of Securin-YFP after activation in control and nocodazole treated oocytes in early metaphase II**

Prophase I oocytes were injected with Securin-YFP mRNA, and were left untreated (Ctrl) or treated with Nocodazole (Noc) for 1h at GVBD+12h. Acquisitions of YFP channel were started 20 min after activation by addition of strontium at GVBD+13h, every 10 min (h: min). Shown are overlays from stack of 15 z-sections spaced by 3 µm. This is a representative movie of 20 control and 17 nocodazole treated oocytes. Related to Figures 2B and S1A.

**Video 3: Visualization of Securin-YFP after activation in control and nocodazole treated oocyte in late metaphase II**

Prophase I oocytes were injected with Securin-YFP mRNA, and were left untreated (Ctrl) or treated with Nocodazole (Noc), for 1h at GVBD+18h. Acquisitions of YFP channel were started 20 min after activation by addition of strontium at GVBD+19h, every 10 min (h: min). Shown are overlays from stack of 15 z-sections spaced by 3 µm. This is a representative movie from 16 control and 10 nocodazole treated oocytes. Related to Figures 2B and S1B.

**Video 4: Super-resolution imaging of the spindle in fixed control and STLC treated oocytes in early metaphase II after *in vitro* maturation**

*In vitro* matured oocytes were let untreated (Ctrl) or treated with STLC at 12 h after GVBD, for 1h, fixed and stained with DAPI (blue), ACA (red) and anti-Tubulin antibody (green). Z-stacks spaced by 0,1 µm were acquired with SIM. Related to Figure S2A-D.

**Video 5: Visualization of anaphase II after activation in control, STLC, and Reversine+STLC treated oocytes in early metaphase II**

Prophase I oocytes were injected with H2B-RFP mRNA, and were left untreated (Ctrl) or treated with STLC or Reversine+STLC, for 1 h at GVBD+12h. Acquisitions of RFP channel were started 20 min after activation by addition of strontium at GVBD+13h, every 10 min (h: min). Shown are overlays from stack of 15 z-sections spaced by 3 µm. This is a representative movie from 35 control, 26 STLC and 37 Reversine+STLC treated oocytes. Related to Figures 2E and S3A.

**Video 6: Visualization of Securin-YFP after activation in control, STLC, and AZD1152+STLC treated oocytes in early metaphase II**

Prophase I oocytes were injected with Securin-YFP mRNA, and were left untreated (Ctrl) or treated with STLC, or AZD1152+STLC, for 1 h at GVBD+12h. Acquisitions of YFP channel were started 20 min after activation by addition of strontium at GVBD+13h, every 10 min (h: min). Shown are overlays from stack of 15 z-sections spaced by 3 µm. This is a representative movie from 17 control, 13 STLC, and 17 AZD1152+STLC treated oocytes. Related to Figures 2F and S3B.

**Video 7: Visualization of anaphase II after activation in control, STLC, and MLN8237+STLC treated oocytes in early metaphase II**

Oocytes incubated with SirDNA, were left untreated (Ctrl) or treated with STLC or MLN8237+STLC, for 1 h at GVBD+12h. Acquisitions of RFP channel were started 20 min after activation by addition of strontium at GVBD+13h, every 10 min (h: min). Shown are overlays from stack of 15 z-sections spaced by 3 µm. This is a representative movie from 29 control, 26 STLC and 27 MLN8237+STLC treated oocytes. Related to Figures 2G and S3C.

**Video 8: Super-resolution imaging of the spindle in fixed control and STLC treated oocytes in late metaphase II after *in vitro* maturation**

*In vitro* matured oocytes were left untreated (Ctrl) or treated with STLC at 18 h after GVBD, for 1h, fixed and stained with DAPI (blue), ACA (red) and anti-Tubulin antibody (green). Z-stacks spaced by 0.1 µm were acquired with SIM. Related to Figure S2E-H.

**Video 9: Visualization of anaphase II after activation in control and STLC treated oocytes in late metaphase II**

Prophase I oocytes were injected with H2B-RFP mRNA, and were left untreated (Ctrl) or treated with STLC at GVBD+18h. Acquisitions of RFP channel were started 20 min after activation by addition of strontium at GVBD+19h, every 10 min (h: min). Shown are overlays from stack of 15 z-sections spaced by 3 µm. This is a representative movie from 41 control and 46 STLC treated oocytes. Related to Figures 2H and S3D.

**Video 10: Visualization of chromosome realignment after STLC wash-out in early metaphase II** Oocytes were incubated with SirDNA and treated with STLC to induce monopolar spindle for 1 h at GVBD+12h. After 1 h, oocytes were washed to remove STLC in the absence (Ctrl) or presence of MLN8237 or AZD1152. Acquisitions of far-red channel were started 10 min after wash-out, every 10 min (h: min). Shown are overlays from stack of 20 z-sections spaced by 3 µm. This is a representative movie from 30 control, 37 MLN8237 and 33 AZD1152 treated oocytes. Related to Figures 4B and 4C.

**Video 11: Visualization of chromosome realignment after STLC wash-out in late metaphase II**

Oocytes were incubated with SirDNA and treated with STLC to induce monopolar spindle for 1 h at GVBD+18h. After 1 h, oocytes were washed to remove STLC in the absence (Ctrl) or presence of MLN8237 or AZD1152. Acquisitions of far-red channel were started 10 min after wash-out, every 10 min (h: min). Shown are overlays from stack of 20 z-sections spaced by 3 µm. This is a representative movie from 35 control, 33 MLN and 37 AZD treated oocytes. Related to Figures 4B and 4D.

**Video 12: Super-resolution imaging of the spindle in fixed control and STLC treated oocytes after *in vivo* maturation 14 h after hCG injection**

*In vivo* matured oocytes collected at 14h after hCG injection, were left untreated (Ctrl) or treated with STLC for 1h, fixed and stained with DAPI (blue), ACA (red) and anti-Tubulin antibody (green). Z-stacks spaced by 0.1 µm were acquired with SIM. Related to Figure 5A-D.

**Video 13: Super-resolution imaging of the spindle in fixed control and STLC treated oocytes after *in vivo* maturation 20 h after hCG injection**

*In vivo* matured oocytes collected at 20h after hCG injection, were left untreated (Ctrl) or treated with STLC for 1h, fixed and stained with DAPI (blue), ACA (red) and anti-Tubulin antibody (green). Z-stacks spaced by 0.1 µm were acquired with SIM. Related to Figure 5A-D.

**Video 14: Visualization of anaphase II after activation of *in vivo* matured oocytes 14 h after hCG injection**

*In vivo* matured oocytes collected at 14h after hCG injection, were incubated with SirDNA and were left untreated (Ctrl) or treated with STLC for 1h. Acquisitions of Far-Red channel were started 20 min after activation by addition of strontium, every 10 min (h: min). Shown are overlays from stack of 20 z-sections spaced by 3 µm. This is a representative movie from 52 control and 43 STLC treated oocytes. Related to Figures 5E.

**Video 15: Visualization of anaphase II after activation of *in vivo* matured oocytes 20 h after hCG injection**

*In vivo* matured oocytes collected at 20h after hCG injection, were incubated with SirDNA and were left untreated (Ctrl) or treated with STLC (1 or 2 µM) for 1h. Acquisitions of Far-Red channel were started 20 min after activation by addition of strontium, every 10 min (h: min). Shown are overlays from stack of 20 z-sections spaced by 3 µm. This is a representative movie from 52 control, 51 STLC 1 µM, and 46 STLC 2µM treated oocytes. Related to Figures 5E.

